# Rotavirus Calcium Dysregulation Manifests as Dynamic Calcium Signaling in the Cytoplasm and Endoplasmic Reticulum

**DOI:** 10.1101/627877

**Authors:** Alexandra L. Chang-Graham, Jacob L. Perry, Alicia C. Strtak, Nina K. Ramachandran, Jeanette M. Criglar, Asha A. Philip, John T. Patton, Mary K. Estes, Joseph M. Hyser

**Affiliations:** Alkek Center for Metagenomic and Microbiome Research; Department of Molecular Virology and Microbiology; Department of Medicine, Gastroenterology and Hepatology, Baylor College of Medicine, Houston, TX 77303; Department of Biology, Indiana University, Bloomington, IN 47405

## Abstract

Like many viruses, rotavirus (RV) dysregulates calcium homeostasis by elevating cytosolic calcium ([Ca^2+^]cyt) and decreasing endoplasmic reticulum (ER) stores. While an overall, monophasic increase in [Ca^2+^]cyt during RV infection has been shown, the nature of the RV-induced aberrant calcium signals and how they manifest over time at the single-cell level have not been characterized. Thus, we generated cell lines and human intestinal enteroids (HIEs) stably expressing cytosolic and/or ER-targeted genetically-encoded calcium indicators to characterize calcium signaling throughout RV infection by time-lapse imaging. We found that RV induces highly dynamic [Ca^2+^]cyt signaling that manifest as hundreds of discrete [Ca^2+^]cyt spikes, which increase during peak infection. Knockdown of nonstructural protein 4 (NSP4) attenuates the [Ca^2+^]cyt spikes, consistent with its role in dysregulating calcium homeostasis. RV-induced [Ca^2+^]cyt spikes were primarily from ER calcium release and were attenuated by inhibiting the store-operated calcium entry (SOCE) channel Orai1. RV-infected HIEs also exhibited prominent [Ca^2+^]cyt spikes that were attenuated by inhibiting SOCE, underlining the relevance of these [Ca^2+^]cyt spikes to gastrointestinal physiology and role of SOCE in RV pathophysiology. Thus, our discovery that RV increases [Ca^2+^]cyt by dynamic Ca^2+^ signaling, establishes a new, paradigm-shifting understanding of the spatial and temporal complexity of virus-induced Ca^2+^ signaling.

## Introduction

Eukaryotic signal transduction pathways employ a variety of signaling molecules to regulate cellular processes. Calcium (Ca^2+^) is one of the most ubiquitous secondary messengers in the cell, which tightly regulates Ca^2+^ movement through the coordinated function of Ca^2+^ channels, transporters, and pumps. Since Ca^2+^ signaling modulates a wide array of cellular processes, it is not surprising that many different viruses exploit Ca^2+^ signaling to facilitate their replication, and the resulting dysregulation of Ca^2+^ signaling causes pathogenesis. Rotavirus (RV), a member of the *Reoviridae* family, is one of the first viruses shown to elevate cellular Ca^2+^ levels and has become a widely-used model system to characterize mechanisms by which viruses dysregulate host Ca^2+^ homeostasis^1^. RV is a clinically important enteric virus that causes severe diarrhea and vomiting in children, resulting in over approximately 258 million diarrhea episodes and 198,000 deaths in 2016^2^. Hyperactivation of cyclic nucleotide (*e.g.*, cAMP/cGMP) and Ca^2+^ signaling pathways is a common strategy among enteric pathogens^3^. Thus, understanding how RV exploits Ca^2+^ signaling is key to understanding and combating RV-induced diarrhea.

RV was first reported to elevate cytosolic [Ca^2+^] by Michelangeli et al. (1991), which stimulated subsequent research into how RV alters cellular Ca^2+^ levels^4^. RV causes a 2-fold steady-state increase in cytosolic Ca^2+^, which is due to increased Ca^2+^ release from the endoplasmic reticulum (ER) and increased Ca^2+^ influx through host Ca^2+^ channels in the plasma membrane (PM)^1, 5^. Elevated cytosolic Ca^2+^ activates autophagy, which is critical for RV replication, and has wide-ranging consequences to host cell functions, including disruption of the cytoskeleton and activation of chloride and serotonin secretion to cause diarrhea and vomiting^1, 5^.

RV dysregulates Ca^2+^ homeostasis by at least two functions of its nonstructural protein 4 (NSP4), a glycoprotein with multiple functions during the infection^5^. In RV-infected cells, ER-localized NSP4 is a viroporin (*i.e.*, virus-encoded ion channel) that releases ER Ca^2+^ and reduces ER Ca^2+^ stores causing a persistent increase in cytosolic Ca^2+^ ^6–8^. Using patch clamp electrophysiology, we demonstrated that the NSP4 viroporin domain (aa47-90) forms a Ca^2+^-permeable ion channel, confirming that NSP4 can directly mediate loss of ER Ca^2+^ ^9^. Decreased ER Ca^2+^ activates stromal interaction molecule 1 (STIM1), an ER Ca^2+^ sensor, which in turn activates Ca^2+^ influx through store-operated calcium entry (SOCE) channels in the PM, primarily Orai1^10^. Voltage-gated Ca^2+^ channels and the sodium-calcium exchanger (NCX) have also been implicated in Ca^2+^ influx in RV-infected cells^11, 12^. Finally, through a mechanistically distinct pathway, RV-infected cells secrete an extracellular NSP4 (eNSP4) cleavage product that elicits a receptor-mediated, inositol triphosphate (IP_3_)-dependent Ca^2+^ signal ^13, 14^. eNSP4 induces diarrhea in neonatal mice, making this the first identified viral enterotoxin ^13, 15^. At the cellular level, the eNSP4-induced Ca^2+^ signal activates chloride secretion through Ca^2+^-activated chloride channels (*e.g.*, anoctamin 1), consistent with its enterotoxin activity *in vivo* ^16, 17^. Thus, NSP4 dysregulates Ca^2+^ by both directly releasing ER Ca^2+^ and by exploiting host SOCE and agonist-induced Ca^2+^ signaling pathways^1^.

While the global dysregulation of Ca^2+^ homeostasis by RV has been well characterized, many mechanistic details about RV-induced Ca^2+^ signaling remain unknown. First, cell population-based studies show that RV induces a monophasic 2-fold increase in cytosolic Ca^2+^ during infection^18, 19^, but whether individual cells manifest this as a monophasic Ca^2+^ increase or a series of discrete Ca^2+^ signals remains unknown. This is an important distinction because the amplitude, duration, and degree of oscillation of cytosolic Ca^2+^ signals regulate downstream pathways. For example, a sustained increase in cytosolic Ca^2+^ activates apoptotic programs, whereas transient or oscillating Ca^2+^ signals activate proliferation and pro-survival pathways^20, 21^. Second, studies in single RV-infected cells have focused on times late post-infection, 7-8 hours post-infection (hpi)^7, 11^. Thus, it is not known whether RV dysregulates Ca^2+^ signaling early in infection or how dysregulation of Ca^2+^ homeostasis progresses during the infection. Finally, RV-induced depletion of ER Ca^2+^ remains controversial due to conflicting data. RV decreases agonist-induced release of ER Ca^2+^ and induces STIM1 activation, suggesting decreased ER Ca^2+^ levels^10, 22^. However, greater uptake of radioactive Ca^2+^ into ER stores has also been observed^7^. As with cytosolic Ca^2+^, ER Ca^2+^ levels are dynamic; yet RV-induced changes to ER Ca^2+^ and how this relates to cytosolic Ca^2+^ signaling remain incompletely characterized.

Addressing these gaps-in-knowledge requires the ability to measure changes in cytosolic and ER Ca^2+^ with single-cell resolution over many hours. For many years, technical limitations of Ca^2+^ indicator dyes (*e.g.*, photobleaching, uneven loading, and dye leakage) made single-cell measurements of Ca^2+^ signaling throughout the entire RV infection not feasible. However, we recently developed the use of genetically-encoded calcium indicators (GECIs) for the study of Ca^2+^ signaling in virus-infected cells ^23^. GECIs are dynamic fluorescent protein-based Ca^2+^ sensors, and variants have been developed for simultaneous Ca^2+^ measurements in multiple cellular compartments (*e.g.*, cytoplasm and ER) ^24^. The stability and targetability of GECIs provides the spatial and temporal resolution needed to perform long-term live-cell Ca^2+^ imaging such that Ca^2+^ signaling can be measured throughout a RV infection. The goal of this study was to use cell lines and human intestinal enteroids (HIEs) stably expressing cytoplasmic and/or ER-localized GECIs to define RV-induced Ca^2+^ signaling dynamics at the single-cell level, and thereby gain new mechanistic insights into how RV dysregulates Ca^2+^ homeostasis.

## Materials and Methods

### Cells and rotaviruses

MA104 cells (African green monkey kidney cells) and HEK293T cells (ATCC CRL-3216) were cultured in high glucose DMEM supplemented with 10% fetal bovine serum (FBS) and Antibiotic/Antimycotic (Invitrogen) at 37°C in 5% CO_2_. Rotavirus SA114F was produced from in-house stocks. Porcine strains OSUv and OSUa were provided as a kind gift from Dr. Lennart Svennson ^25^, and the human strain Ito was prepared as previously described ^26^. Recombinant SA11 clone 3 expressing an mRuby3 red fluorescent protein reporter downstream of NSP3 (SA11-mRuby) was generated using a modified plasmid based reverse genetics (RG) system ^27^. Briefly, the NSP4 open reading frame (ORF) in the pt7/NSP3 plasmid was replated with an ORF encoding NSP3 fused downstream to FLAG-tagged mRuby3. To promote the translation of NSP3 and mRuby3 as separate proteins, we inserted a teschovirus 2A-like stop-restart translation element between the NSP3 and FLAG-tagged mRuby3 coding sequences^28^. The SA11-mRuby virus was generated by co-transfection of BHK-T7 cells with pT7 plasmids expressing RV plus-sense RNAs along with a plasmid expressing the African swine fever virus NP868R capping enzyme from a CMV promoter. All viruses were propagated in MA104 cells in serum-free DMEM supplemented with 1 μg/mL Worthington’s Trypsin, and after harvest stocks were subjected to three freeze/thaw cycles and activated with 10 μg/mL Worthington’s Trypsin for 30 min at 37 °C prior to use.

### Chemicals

2-APB, KB-R7943 mesylate, BTP2 (YM 58483), and BAPTA-AM were purchased from Tocris Bioscience. Methoxyverapamil (D600), nitrotetrazolium blue (NBT), and BCIP were purchased from Sigma-Aldrich. Synta66 and GSK7975A were purchased form Aobious. EGTA solution (0.5 M, pH 8.0) was purchased from Invitrogen.

### Antibodies

To detect RV, we used rabbit anti-RV strain Alabama ^29^ (IF, 1:80,000; western blot, 1:3000), guinea pig anti-NSP2 ^30^ (IF, 1:5000), and rabbit anti-NSP4 aa120-147 ^31^ (western blot, 1:3000). Secondary antibodies for IF were donkey anti-rabbit AlexaFluor 568 (Invitrogen) and donkey anti-guinea pig Dylight 549 (Rockland), both at 1:2000. For western blots, we used mouse anti-GAPDH monoclonal antibody (Lifetein) (1:5000) and secondary antibodies alkaline phosphatase-conjugated goat anti-rabbit IgG or goat anti-mouse IgG (Southern Biotech) (1:2000).

### Calcium indicator lentiviruses, cell lines, and enteroids

GCaMP5G (Addgene plasmid #31788), GCaMP6s (Addgene plasmid #40753), and G-CEPIA1er (Addgene plasmid #105012) were cloned into pLVX-Puro. RGECO1.2 (Addgene plasmid #45494), R-CEPIA1er (Addgene plasmid #58216), and GCaMP6s were cloned into pLVX-IRES-Hygro. Lentivirus vectors for the GECI constructs were packaged in HEK293T cells as previously described ^23^ or produced commercially (Cyagen Biosciences, Inc.).

Production of the MA104-GCaMP5G and MA104-GCaMP5G/RCEPIAer cell lines were previously described and similar methods were used to generate MA104-RGECO1/GCEPIAer and the MA104-GCaMP6s-shRNA lines ^23^. Human intestinal enteroids expressing GCaMP6s (G6S-HIEs) were created using lentivirus transduction as described previously and grown in high Wnt3a CMGF+ with 1 µg/mL puromycin for selection^32^. Proper GECI functionality was validated by responses to 50 μM ATP.

### MA104-GCaMP6s cells expressing NSP4 shRNAs

Lentivirus constructs encoding shRNA targeting SA11 gene 10 and a non-targeting scrambled shRNA negative control were generated and packaged by America Pharma Source, LLC. The NSP4-shRNA1 targets gene 10 nt50-70 (5’-GCTTACCGACCTCAATTATAC-3’) and NSP4-shRNA2 targets gene 10 nt176-196 (5’-GCTACATAAAGCATCCATTCC-3’). The shRNA-expression vectors encode a blasticidin-resistance gene for drug selection. Parental MA104 cells were transduced with the shRNA-expressing constructs and at 72 hrs post-transduction the cells were passaged in the presence of 30 µg/mL blasticidin to select for stably transduced cells. These three cell lines were then transduced with GCaMP6s GECI (in pLVX-IRES-Hygro) and passaged with 50 μg/mL hygromycin B and 30 μg/mL blasticidin to select for co-expression of GCaMP6s and the shRNA.

### Establishment of HIE cultures

Three-dimensional human intestinal enteroid (HIE) cultures were generated from crypts isolated from adult patients undergoing bariatric surgery as previously described^26, 33^. These established cultures were obtained at Baylor College of Medicine through the Texas Medical Center Digestive Diseases Center Gastrointestinal Experimental Model Systems Core. For these studies, jejunum HIEs from patient J3 were used. Complete media with and without growth factors (CMGF+ and CMGF-, respectively), differentiation media, and high Wnt3a CMGF+ (hW-CMGF+) were prepared as previously described^26, 33, 34^. Fluorobrite DMEM supplemented with 15mM HEPES, 1X sodium pyruvate,1X Glutamax, and 1X non-essential amino acids (Invitrogen) was used for fluorescence Ca^2+^ imaging (FB-Plus). An FB-Plus-based differentiation medium (FB-Diff) consisted of FB-Plus with the same added components as differentiation media, but without Noggin. HIEs were grown in phenol red-free, growth factor-reduced Matrigel (Corning). G6S-jHIE monolayers were prepared from three-dimensional cultures and seeded into optical-bottom 10-well Cellview chamber slides coated with dilute collagen IV (Sigma) as described previously^35, 36^. After 24 hr in CMGF+ and 10 µM Y-27632 Rock inhibitor, differentiation medium was used and changed every day for 4-5 days.

### Microscopy and image analysis

To image viroplasms, we used a GE Healthcare DeltaVision LIVE High Resolution Deconvolution with an Olympus IX-71 base and illumination provided by a xenon lamp. Images were captured with Plan Apo 60X Oil DIC objective and a pco.edge sCMOS camera. Images were acquired and deconvolved using SoftWoRx software and further analyzed with Fiji (ImageJ).

For calcium imaging, MA104 cells and HIEs were imaged with a widefield epifluorescence Nikon TiE inverted microscope using a SPECTRAX LED light source (Lumencor) and either a 20x Plan Fluor (NA 0.45) phase contrast or a 20X Plan Apo (NA 0.75) differential interference contrast (DIC) objective. Fluorescence and transmitted light images were recorded using an ORCA-Flash 4.0 sCMOS camera (Hamamatsu), and Nikon Elements Advanced Research v4.5 software was used for multipoint position selection, data acquisition, and image analysis.

Images were read-noise subtracted using an average of 10 no-light acquisitions of the camera. Single cells were selected as Regions of Interest (ROI) and fluorescence intensity measured for the experiment. 3D HIE’s fluorescence was measured individually using threshold analysis adjusted to select each enteroid with the Fill Holes algorithm included. Enteroids that moved out of the field of view or could not be separated from adjacent enteroids were removed from analysis. The fluorescence intensity of whole field-of-view was measured for HIE monolayers.

Fluorescence intensity values were exported to Microsoft Excel and normalized to the baseline fluorescence. The number and magnitude of Ca^2+^ spikes were calculated by subtracting each normalized fluorescence measurement from the previous measurement to determine the change in GECI fluorescence (ΔF) between each timepoint. Ca^2+^ signals with a ΔF magnitude of > 5% were counted as Ca^2+^ spikes.

### Calcium imaging

#### MA104-GECI cells

Confluent monolayers of MA104-GECI cells in 8-well chamber slides (ibidi) were mock- or RV-infected in FBS-free media for 1 hr at the indicated multiplicity of infection (MOI). Then the inoculum was removed and replaced with FB-Plus, and for appropriate studies, with DMSO or drugs at indicated concentrations. The slide was mounted into an Okolab stage-top incubation chamber equilibrated to 37°C with a humidified 5% CO_2_ atmosphere. For each experiment, 3-5 positions per well were selected and imaged every 1 minute for ∼18-20 hrs.

#### GECI HIEs

To test Ca^2+^ response, 3D G6S-jHIEs were suspended in 25% Matrigel diluted in FB-Diff media and seeded into optical-bottom 10-well Cellview chamber slides (Greiner bio-one) thinly coated with Matrigel. After baseline imaging using the stage-top incubator, 200µM carbachol in FB-Diff or FB-Diff alone was added to the well and imaging continued for 1 hour with 6-10 enteroids imaged every 10 s.

For RV infection in 3D HIEs, the jHIEs were split and grown in hW-CMGF+ for 2 days followed by differentiation medium for 1 day. G6S-jHIEs were gently washed using ice cold 1XPBS and resuspended in inoculum of 50µL MA104 cell lysate or RV (strain Ito) diluted with 150 µL CMGF- and incubated for 1 hr. Then HIEs were washed, resuspended in 25% Matrigel diluted in FB-Diff (with DMSO or 2-APB in indicated experiments) and pipetted onto 8-well chamber slides (Matek) pre-coated with Matrigel. Imaging positions were chosen so that between 20-50 enteroids were selected per experimental condition. Enteroids were imaged using the stage-top incubator with transmitted light and GFP fluorescence every 2-3 minutes for ∼18 hrs.

For RV infection in monolayers, G6S-jHIE monolayers were washed once with CMGF- and treated with an inoculum of 50µL CMGF-plus 30µL MA104 cell lysate or RV (strain Ito) and incubated for 2 hr. Then inoculum was removed, and monolayers were washed once with FB-Diff before adding FB-Diff with DMSO or 2-APB. Monolayers were transferred to the stage-top incubator for imaging with 4 fields of view chosen per well, and GFP fluorescence was measured every minute for ∼18 hrs.

### Store-operated calcium entry assay

G6S-jHIE monolayers after 4 days in differentiation media were washed and incubated in 0 mM Ca^2+^ (0Ca^2+^) Ringers solution (160 mM NaCl, 4.5 mM KCl, 1 mM MgCl_2_, 10 mM HEPES, pH=7.4). Endoplasmic reticulum Ca^2+^ stores were depleted by incubating cells with 500 nM thapsigargin in 0Ca^2+^ with either 50 µM 2-APB or DMSO as a vehicle control. SOCE was measured using live-cell fluorescence imaging of the increase in GFP fluorescence after the addition of normal Ringers to bring the total Ca^2+^ concentration to 2 mM.

### Western blot analysis

RV proteins were detected by immunoblot analysis as previously described, with the following modifications^31^. Cells were lysed using a 1X RIPA buffer solution [10 mM Tris-HCL pH 8.0, 1 mM EDTA, 1% Triton X-100, 0.1% sodium deoxycholate, 0.1% SDS, 140 mM NaCl, and 1 tablet complete mini protease inhibitor (Roche)] and passed through a Qiashredder (Qiagen). Samples were boiled for 10 min at 100°C in SDS-PAGE sample buffer and separated on Tris-glycine 4-20% SDS-PAGE gels (BioRad). Detected protein bands for each blot were quantified using ImageJ software for gel densitometry measurements of NSP4:GAPDH.

### Immunofluorescence

MA104 cells and HIEs were fixed using the Cytofix/Cytoperm kit (BD Biosciences) according to manufacturer instructions. Primary antibodies were diluted in 1X Perm/Wash overnight at 4°C. The next day, the cells were washed three times with 1X Perm/Wash solution and then incubated with corresponding secondary antibodies for 1 hr at room temperature. Nuclei were stained with NucBlue Fixed Cell Stain (Life Technologies) for 5 min at room temperature and washed with 1X PBS for imaging.

### Plaque assays

Plaque assays were performed as described previously with the following modifications ^37^. Briefly, MA104 cells or the MA104-shRNA expressing cells were seeded and grown to confluency in 6 wells plates. Wells were infected at 10-fold dilutions in duplicate for 1 hr and media replaced with an overlay of 1.2% Avicel in serum-free DMEM supplemented with DEAE dextran, and 1μg/mL Worthington’s Trypsin, and, for indicated experiments, DMSO vehicle or SOCE drugs^38^. The cells were incubated at 37°C/5% CO_2_ for 48-72 hrs before overlay was removed and cells stained with crystal violet to count plaques.

### RNA extraction, reverse transcription, and quantitative PCR

Total RNA was extracted from HIEs wells (in hW-CMGF+ or differentiation media for 4 days) or MA104 cells grown to confluency in a 6-well plate using TRIzol reagent (Ambion). Total RNA was treated with Turbo DNase I (Ambion) and cDNA was generated from 250 ng RNA using the SensiFAST cDNA synthesis kit (Bioline). Quantitative PCR was performed using Fast SYBR Green (Life Technologies) with primers designed using NCBI Primer-Blast (Table 1) and using a QuantStudio real time thermocycler (Applied Biosciences). Target genes were normalized to the housekeeping gene ribosomal subunit 18s and relative expression was calculated using the ddCT method.

**Table 1:**
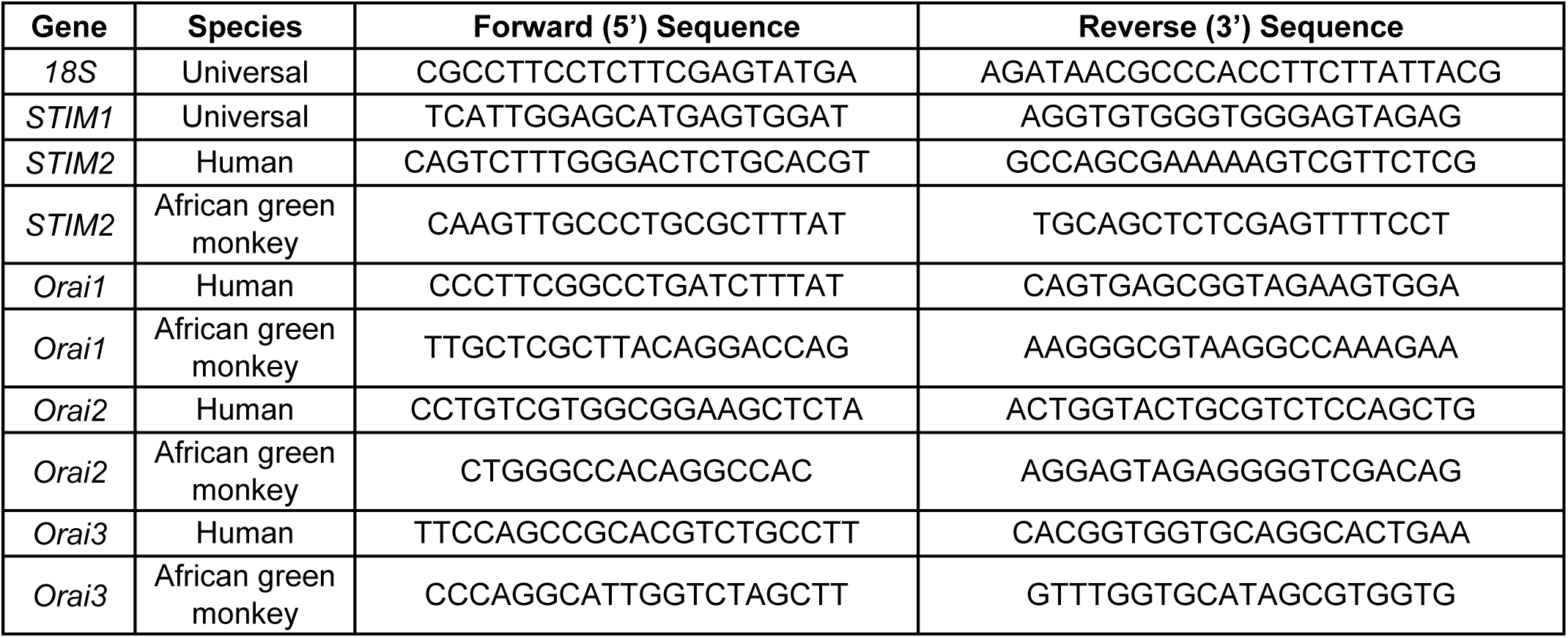
Primers used in this work.

### Statistical analysis

Biostatistical analyses were performed using GraphPad Prism (version 8.1) with results presented as mean ± standard deviation. Comparisons used an unpaired Student’s t-test, the nonparametric Mann-Whitney test, or a One-way Analysis of Variance (ANOVA) with Tukey’s post hoc multiple comparisons test where appropriate. Differences were considered statistically significant for p < 0.05. All authors had access to the study data, reviewed, and approved the final manuscript.

## Results

Previous studies have shown that RV significantly increases cytosolic Ca^2+^ over several hours during the peak of RV replication^18, 39^; however, the kinetics of this increase and whether it is a monophasic increases or manifests as discrete Ca^2+^ transients are not known. To address these questions, we developed a series of cell lines stably expressing GECIs and used these cell lines to perform live-cell Ca^2+^ imaging over the course of a RV infection^23^. For the long-term imaging experiments, MA104 cells stably expressing cytosolic GCaMP5G (MA104-GCaMP5G) were seeded into chamber slides and either mock- or RV-infected with strain SA114F (MOI 10), and GCaMP5G fluorescence imaged for ∼18 hr (2-20 hpi) (Fig. 1). Mock-infected cells maintained a low fluorescence throughout the time course (Fig. 1A, upper panels), whereas RV-infected cells exhibited strongly increased fluorescence, indicating elevated Ca^2+^ levels (Fig. 1A, lower panels and Supplementary Video 1 online). Since GECIs use an engineered calmodulin to sense Ca^2+^, overexpression of GCaMP5G could act as a Ca^2+^ buffer and alter the kinetics of RV replication. To assess whether the MA104-GCaMP5G cells exhibited altered RV infection/protein synthesis, we analyzed NSP4 expression in parental MA104 cells and the MA104-GCaMP5G cells infected with SA114F (MOI 10) from 3-8 hpi by western blot (Fig. 1B). NSP4 expression was similar in both parental and GCaMP5G-expressing cells, indicating that stable GCaMP5G expression does not interfere with RV infection.

**Figure 1.**
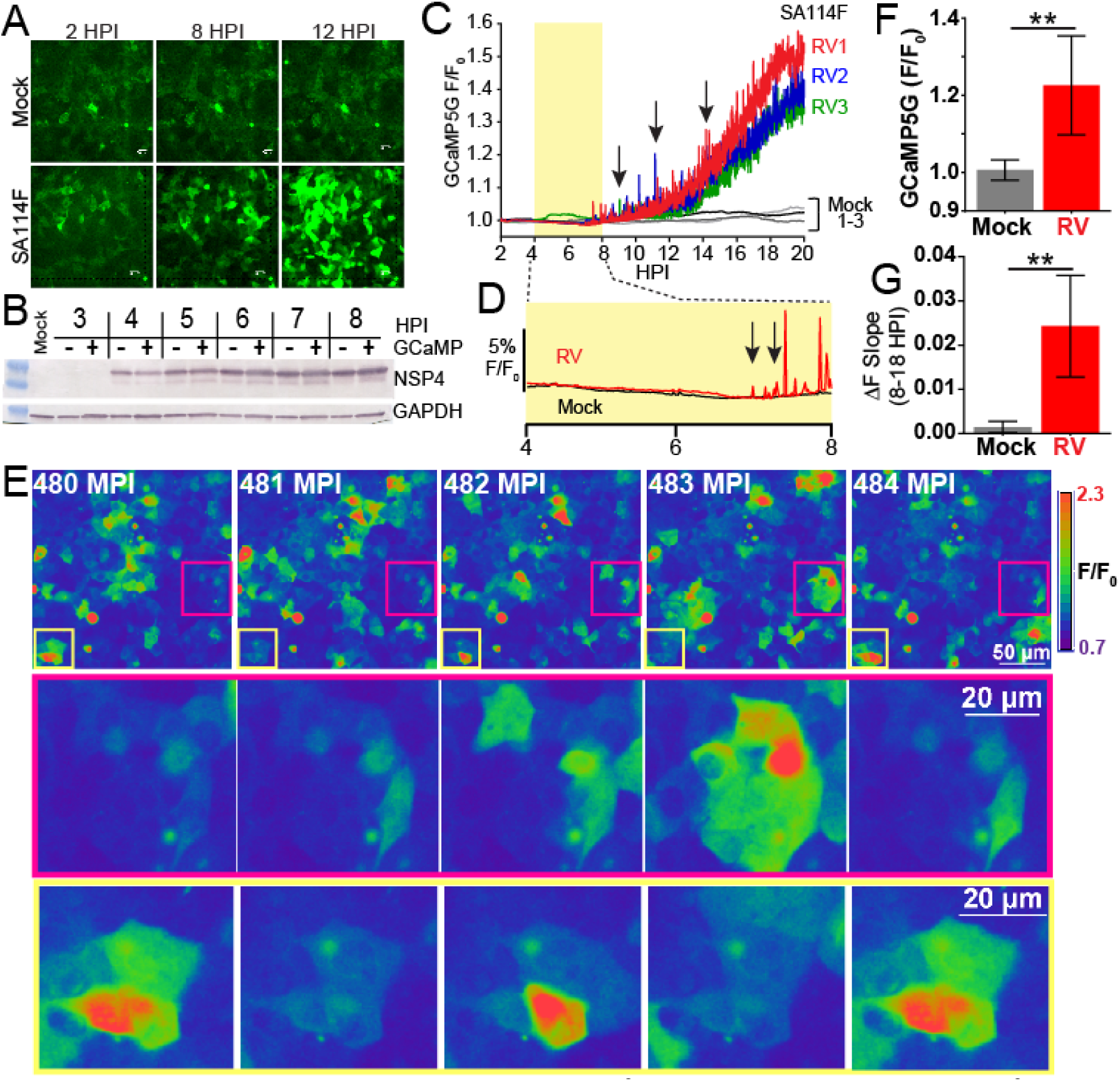
RV-induced increase in cytosolic Ca^2+^ manifests as increased Ca^2+^ signaling dynamics. **(A)** Epifluorescence images of mock (upper) or SA114F-infected (MOI 10) MA104-GCaMP5G cells at 2, 8, and 12 hours post-infection (HPI). **(B)** Western blot of RV NSP4 expression in MA104 cells without (-) or with (+) GCaMP5G from 3-8 HPI. Western blot for GAPDH serves as a loading control. **(C)** GCaMP5G fluorescence (F/F_0_) for three fields-of-view (∼455 µm^2^) each of mock (black and grey lines) or RV-infected (red, blue, and green) MA104-GCaMP5G cells from 2-20 HPI. Increased frequency of transient Ca^2+^ fluxes (arrows) highlight the increased Ca^2+^ signaling dynamics. **(D)** Expanded graph of relative GCaMP5G (F/F_0_) of representative Mock (black) and RV-infected (red) cells. Arrows indicate the increased low/moderate amplitude Ca^2+^ signals present during the initial increase in steady-state cytosolic Ca^2+^. **(E)** Examples of the dynamic Ca^2+^ signaling. GCaMP5G fluorescence pseudocolored by intensity from 480-484 minutes post-infection (MPI). Regions in the magenta and yellow boxes are magnified below. **(F)** Average GCaMP5G fluorescence from 18-19 HPI and **(G)** slope of GCaMP5G ΔF from 8-18 HPI. Data shown as mean ± SD from 12 fields-of-view (triplicate of four independent experiments). **p<0.01

Next, we measured changes in cytosolic Ca^2+^ by determining the relative GCaMP5G fluorescence (F/F_0_) for the whole field-of-view (FOV) (∼455 µm^2^) for three replicate infections and time lapse images were acquired once per minute (Fig. 1C). Mock-infected cells maintained low Ca^2+^ levels with few transient and low amplitude Ca^2+^ signals (Fig. 1C, black and grey lines). In RV-infected cells, the steady-state Ca^2+^ levels began to increase at ∼6 hpi, and we observed many large amplitude, transient Ca^2+^ signals that occurred concomitantly with the steady-state elevation in cytosolic Ca^2+^ levels (Fig. 1C, arrow). Further, at 6-8 hpi the RV-infected cells had more low and moderate amplitude Ca^2+^ signals than mock-infected cells (Fig. 1D, arrows), which occurred during the initial increase in steady-state Ca^2+^ levels. A more detailed examination of the Ca^2+^ signaling over a period of 5 mins at 480 minutes post-infection (∼8 hpi) shows that the increase in cytosolic Ca^2+^ manifests as discrete and dynamic Ca^2+^ fluxes from individual or small groups of cells (Fig. 1E). The dynamic changes are exemplified by the two areas outlined (Fig. 1E, magenta or yellow box), showing substantial changes over the 5 min period. To compare our GECI-based Ca^2+^ imaging of RV-induced Ca^2+^ signaling to previous cell population-based studies, we determined the average GCaMP5G fluorescence from 18-19 hpi (Fig. 1F) and the slope of the fluorescence increase from 8-18 hpi (Fig. 1G). We found a similar ∼2-fold increase in [Ca^2+^]c and a rate of Ca^2+^ increase consistent with that found in studies using Ca^2+^ indicator dyes ^12, 18, 19^. Thus, the MA104-GCaMP5G cells exhibit the well-characterized hallmarks of RV-induced Ca^2+^ dysregulation but have greater spatial and temporal resolution to study Ca^2+^ in RV-infected cells. This has revealed a new dimension of the RV-induced Ca^2+^ signaling, in that the cytosolic Ca^2+^ increase manifests through highly dynamic and discrete Ca^2+^ signaling events, which had not been previously observed.

### RV-induces dynamic Ca^2+^ signaling

Our long-term Ca^2+^ imaging approach using MA104-GCaMP5G cells had sufficient resolution to enable analysis of Ca^2+^ signaling at the single-cell level over the course of the RV infection. Three representative traces from mock- or RV-infected cells (MOI 10, imaged once per minute) show that while individual cells display unique characteristics of Ca^2+^ signaling, they all exhibit similar patterns of Ca^2+^ signaling (Fig. 2A). Mock-infected cells maintain low cytosolic Ca^2+^ with few low amplitude Ca^2+^ signals (Fig. 2A, black lines), but RV-infected cells display a large number of large amplitude Ca^2+^ transients, as well as an overall increase in cytosolic Ca^2+^ (Fig. 2A, red lines). These Ca^2+^ transients were the most prominent Ca^2+^ signal during the infection and were infrequently detected in mock-infected cells. Finally, as RV is a lytic virus, we observed clear evidence of cell lysis late in infection, but this was associated with a loss of GCaMP5G signal and dynamics because the sensor diffused away from the ruptured cells (see Supplementary Video 1 online). We sought to determine a threshold to define these “Ca^2+^ spikes” so that we could measure the number and amplitude of these Ca^2+^ signals. To define a “Ca^2+^ spike”, we set a cutoff for the change in GCaMP5G fluorescence between two measurements to be greater than 5% (ΔF > 5%). The mean Ca^2+^ transient amplitude of mock-infected cells was 0.3% (± 0.5% standard deviation) (data not shown; see Fig. 2D for a subset of this data). Thus the ΔF > 5% cutoff is more than 3 standard deviations above the mean, which establishes a stringent threshold for quantitating Ca^2+^ spikes. Next, we determined the change in fluorescence between each data point and found that the majority of Ca^2+^ spikes were captured in 1 image (Fig. 2B). This simple method enabled detection of Ca^2+^ spikes with approximately 80% accuracy; however, it results in a 20% under-estimation of Ca^2+^ spikes, which were captured in 2-3 images (Fig. 2C, black dots). Nevertheless, using this method of Ca^2+^ spike analysis, we found that RV significantly increases the number of Ca^2+^ spikes per cell (Fig. C). We then determined the amplitude for the top 150 Ca^2+^ spikes per representative cell and found that while the amplitude was highly variable between RV-infected cells, these signals were significantly greater than mock-inoculated cells (Fig. 2D). Finally, we compared the RV-induced Ca^2+^ spikes to the Ca^2+^ response induced by 10 μM ATP (Fig. 2E). As expected, ATP induced a strong Ca^2+^ flux that was similar to the amplitude of Ca^2+^ spikes induced during RV infection. Although the ATP-induced response was significantly greater than the RV-induced Ca^2+^ spikes, it is important to note that the amplitude of the ATP-induced signals represent the peak of the Ca^2+^ response, whereas it is not possible to know how many of the RV-induced Ca^2+^ spikes were captured at the peak of the signal. Thus, at the individual cell level, a high MOI RV infection induces up to hundreds of discrete and high amplitude Ca^2+^ signaling events.

**Figure 2.**
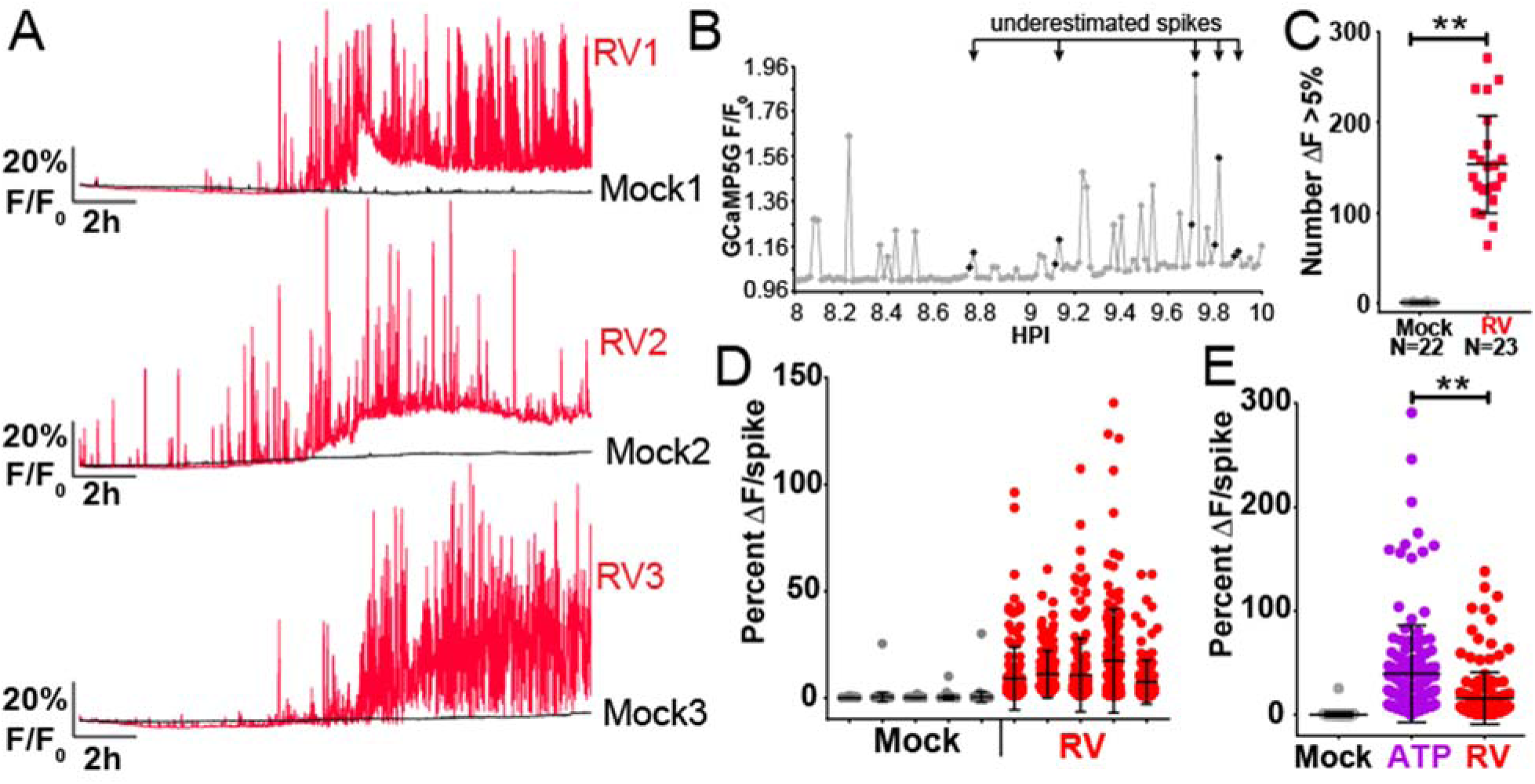
RV increases the number and amplitude of Ca^2+^ spikes at the single-cell level. GCaMP5G (F/F_0_) fluorescence traces from three representative single mock (black) or RV-infected (red) MA104-GCaMP5G cells. **(B)** GCaMP5G fluorescence (F/F_0_) of a single RV-infected MA104-GCaMP5G cell from 8-10 HPI. Symbols in magenta denote Ca^2+^ spikes captured in two data points and potentially underestimated by counting spikes as ΔF >5%. **(C)** MA104-GCaMP5G cells mock or RV infected (MOI 10), GCaMP5G fluorescence was imaged from 2-20 HPI and the number of Ca^2+^ spikes (ΔF >5%) individual cells was determined. **(D)** Ca^2+^ spike amplitude of the top 150 Ca^2+^ spikes from five representative mock and RV-infected cells. **(E)** GCaMP5G Ca^2+^ response to 50 μM ATP in comparison to mock or RV infection. Data shown as mean ± SD. **p<0.01

Next, we determined how these RV-induced Ca^2+^ signals differ with respect to different infectious doses. We infected MA104-GCaMP5G cells with SA114F or with a recombinant SA11 cl. 3 expressing mRuby from the NSP3 gene (SA11cl3-mRuby) ^28^. Cells were infected at MOI of 10, 1, or 0.1, and we performed time-lapse Ca^2+^ imaging and single-cell analysis of the resulting Ca^2+^ signaling. Infection of cells with native SA114F at different MOIs showed the expected infectious dose-dependent increase in the number of RV-positive cells (Fig. 3A). Similarly, the SA11cl3-mRuby-infected cells exhibited an infectious dose-dependent increase in the number of RFP-positive cells at 7 hpi (Fig. 3B), as well as an increase in RFP intensity from 7 hpi to 10 hpi (Fig. 3C). Representative single-cell Ca^2+^ traces for SA114F-infected cells show similar dynamic increases in cytosolic Ca^2+^ spikes as before, but cells infected with lower MOIs of 1 and 0.1 exhibited a later onset of the Ca^2+^ signaling and generally fewer and lower amplitude Ca^2+^ spikes (Fig. 3D). The virus dose-dependent differences in the Ca^2+^ signaling are clearly demonstrated by the time-lapse imaging at 6-7 hpi, which are superimposed onto the immunofluorescence images to detect RV-positive cells (Supplementary Video 2 online). In a more detailed examination of cytosolic Ca^2+^ in RV-infected cells, we used a higher image acquisition frequency (1 image/1.5 sec) and again observed active and dynamic Ca^2+^ signaling. While virtually every cell exhibited multiple Ca^2+^ transients over the course of 10 mins, the Ca^2+^ spike frequency and amplitude were variable from cell-to-cell (Supplemental Figure 1 & Supplementary Video 3 online). We quantitated the number of Ca^2+^ spikes throughout the infection and found a dose-dependent decrease in the number of spikes per cell (Fig. 3F) for lower MOI infections. A similar phenotype was observed in cells infected with SA11cl3-mRuby, but in this case the mRuby expression enabled us to measure both Ca^2+^ signaling and RV protein expression (Fig. 3E & Supplementary Video 4 online). We again observed that for lower MOI infections the onset of Ca^2+^ signaling was later, and the Ca^2+^ spike number and amplitude were generally lower. Further, onset of the Ca^2+^ signaling corresponded to the detection of mRuby from the NSP3 gene. Quantitation of the number of Ca^2+^ spikes per cell for SA11cl3-mRuby infections also showed a dose-dependent decrease with infectious dose (Fig. 3G). Thus, the number and amplitude of these Ca^2+^ signaling events are related to both the infectious dose of RV and onset of RV protein synthesis.

**Figure 3.**
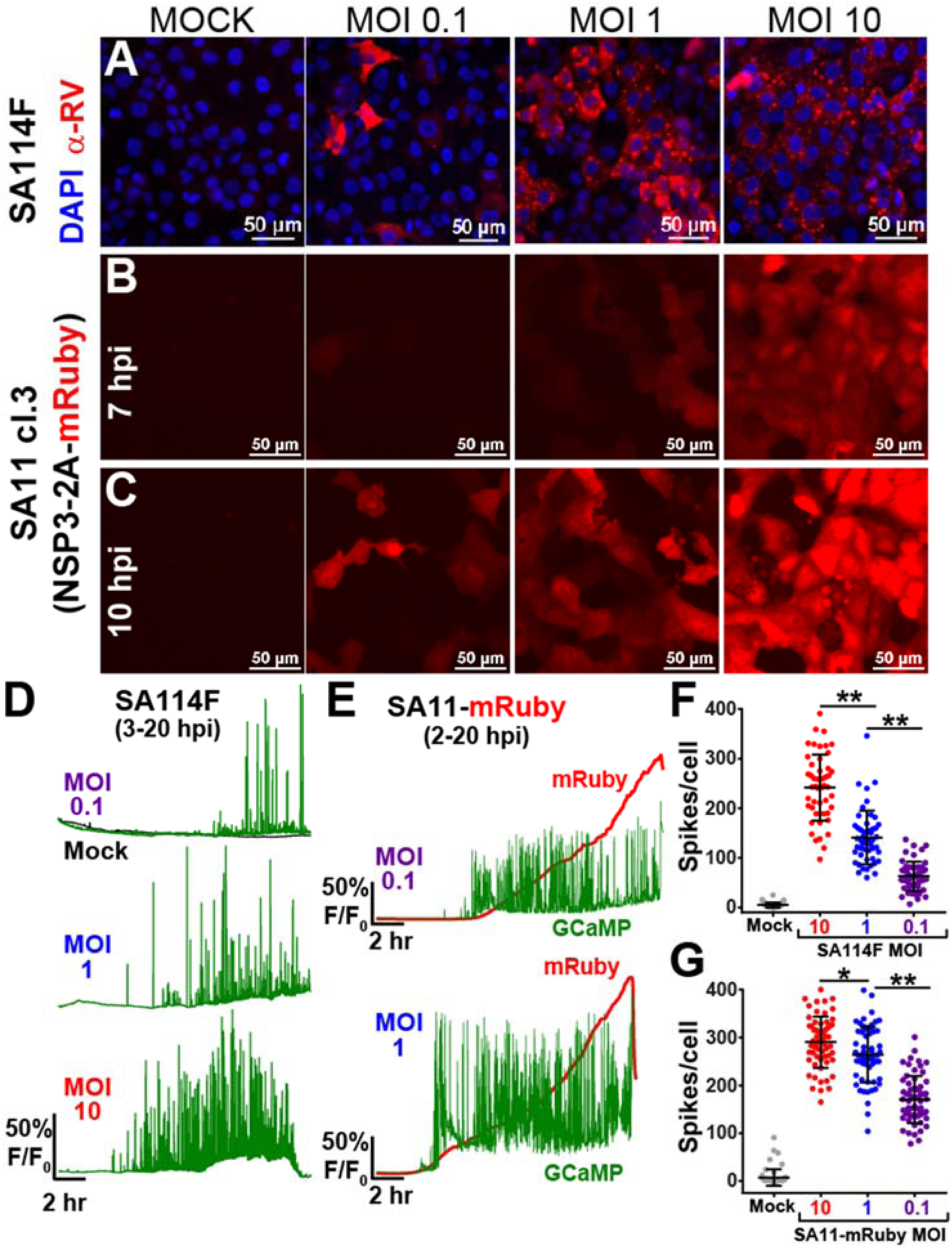
RV-induced dynamic Ca^2+^ signaling is related to virus dose. **(A)** Immunofluorescence images of mock or SA114F-infected MA104-GCaMP5G cells that were inoculated with increasing MOI (0.1, 1, or 10). RV antigen (red) is detected at ∼8 hpi with anti-RV polyclonal antisera and nuclei are stained with DAPI (blue). **(B-C)** Epifluorescence images of MA104-GCaMP5G cells mock or infected with recombinant SA11-mRuby reporter virus with increasing MOIs (0.1, 1, 10). Images were captured at 7 HPI **(B)** or 10 HPI **(C)**. **(D)** Representative single-cell traces of relative GCaMP5G fluorescence (F/F_0_) from cells mock (black) or RV infected by SA114F with MOIs of 0.1 (purple), 1 (blue), 10 (red). **(E)** Representative single-cell traces of relative fluorescence (F/F_0_) of GCaMP5G (green) and mRuby (red) from cells infected by SA11-mRuby MOI 0.1 (purple) or 1 (blue). **(F-G)** Number of Ca^2+^ spikes (F/F_0_ > 5%) from mock or RV-infected cells that were infected with SA114F **(F)** or SA11-mRuby **(G)**. Data shown as mean ± SD of 60 cells/condition. **p<0.01

Next, we sought to characterize the Ca^2+^ signaling phenotype of different RV strains that infect humans or other animals. We compared the Ca^2+^ signaling in MA104-GCaMP5G cells infected at MOI 1 with simian strain SA114F to that of human strain Ito. Immunofluorescence staining of Ito-infected cells (Fig. 4A) showed a similar number of infected cells as that for SA114F infected above (Fig. 3A). As above, we quantitated the number of Ca^2+^ spikes per cell and found that both SA114F and Ito induced a significant increase in Ca^2+^ spikes compared to mock-infected cells (Fig. 4B). These findings demonstrate that the dynamic Ca^2+^ signaling phenotype is not exclusively a feature of animal RV strains. Further, we investigated the attenuated and virulent porcine OSU strains (OSUa and OSUv). Mutations in the OSUa NSP4 protein are associated with reduced elevation in cytosolic Ca^2+^ levels in recombinant NSP4-expressing Sf9 cells ^40^, but the Ca^2+^ signaling phenotype of these two viruses had not been studied in the context of an infection. Immunofluorescence of OSUa and OSUv-infected MA104 cells at MOI 1 show a similar number of infected cells (Fig. 4A). However, while both OSUa and OSUv significantly increase the number of Ca^2+^ spikes, the number of Ca^2+^ spikes from OSUa-infected cells is significantly less than that of those infected with OSUv (Fig. 4B & Supplementary Video 5 online). To characterize this difference further, we examined single-cell traces for OSUa- and OSUv-infected cells (Fig. 4C). We found that OSUa-infected cells initially induced a low-amplitude monophasic increase in cytosolic Ca^2+^ levels (Fig. 4C, black arrows), with the onset of the dynamic Ca^2+^ spikes occurring several hours later (Fig. 4C, purple traces). In contrast, OSUv infection induced a much earlier onset of the dynamic Ca^2+^ spiking, which explains the higher number of Ca^2+^ spikes per cell (Fig. 4C, blue traces). Interestingly, OSUv may also induce an early, low-amplitude increase in cytosolic Ca^2+^ in addition to the dynamic Ca^2+^ spikes, but the high number of Ca^2+^ spikes makes it difficult to clearly ascertain this in all but a few cells (Fig. 4C, red arrow). Together, these data demonstrate that the dynamic Ca^2+^ signaling phenotype is a commonly feature among RV strains and potentially related to NSP4’s role in dysregulating host Ca^2+^ homeostasis and virus virulence.

**Figure 4.**
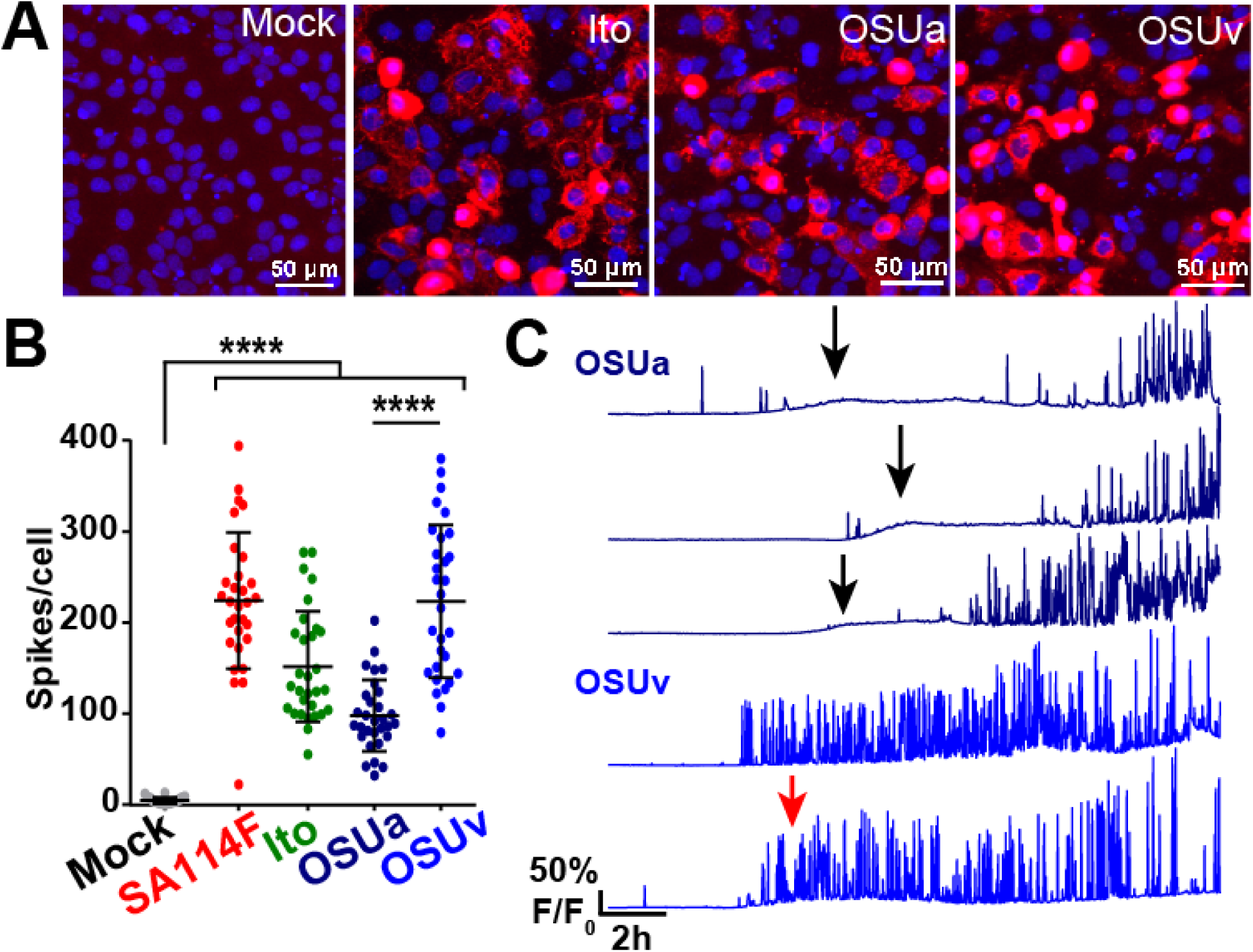
Dynamic Ca^2+^ signaling is induced by several other RV strains. **(A)** Immunofluorescence images of MA104-GCaMP5G cells mock or infected by human RV strain Ito, porcine OSUa, or porcine OSUv at MOI 1. RV antigen (red) is detected at ∼8 hpi with anti-RV polyclonal antisera and nuclei are stained with DAPI (blue). **(B)** Number of Ca^2+^ spikes (F/F_0_ > 5%) from mock or RV-infected cells inoculated with MOI1 of the strains listed. Data shown as mean ± SD of 30 cells/condition. ****p<0.0001. **(C)** Representative single-cell traces of relative GCaMP5G fluorescence (F/F_0_) from cells infected by OSUa (purple) or OSUv (blue). A slight increase in the steady-state Ca^2+^ level is exhibited by most OSUa-infected (black arrows) and some OSUv-infected (red arrow) cells.

### Ca^2+^ signaling dynamics are dependent on NSP4 expression

RV NSP4 is the primary mediator of elevated Ca^2+^ levels during RV infection. The differences in Ca^2+^ signaling by OSUa and OSUv observed in Fig. 4 suggest that NSP4 expression is important for the induction of the dynamic Ca^2+^ signaling during infection^1^. To test the role of NSP4 in these Ca^2+^ signals, we made two GCaMP6s cell lines, each stably expressing a different short-hairpin RNA that targets SA11 NSP4 (NSP4 shRNA1 and NSP4 shRNA2), and a third GCaMP6s cell line stably expressing a non-targeted scrambled shRNA. Cells were infected with SA114F (MOI 0.01), and we found that cells expressing NSP4-targeted shRNAs exhibited knockdown of NSP4 protein levels (Fig. 5A). We normalized NSP4 expression to GAPDH and found ∼40% knockdown in cells expressing NSP4 shRNA1 and ∼85% knockdown in cells expressing NSP4 shRNA2 (Fig. 5B), which correlated with reduced RV plaque size (Fig. 5C). To examine the Ca^2+^ signaling phenotype, we then infected the cells with SA114F (MOI 0.1) and used live-cell imaging to measure Ca^2+^ signaling from ∼2-18 hpi. The scrambled shRNA-expressing cells exhibited a similar degree of dynamic Ca^2+^ signaling as observed in parental MA104 cells (Fig. 5D, red traces), whereas knockdown of NSP4 substantially decreased the degree of Ca^2+^ signaling observed (Fig. 5D, blue traces). Upon quantitation, we found the number of Ca^2+^ spikes was significantly reduced in the NSP4 knockdown cells (Fig. 5E). Together, these data show that NSP4 is responsible for inducing these dynamic Ca^2+^ signals during infection.

**Figure 5.**
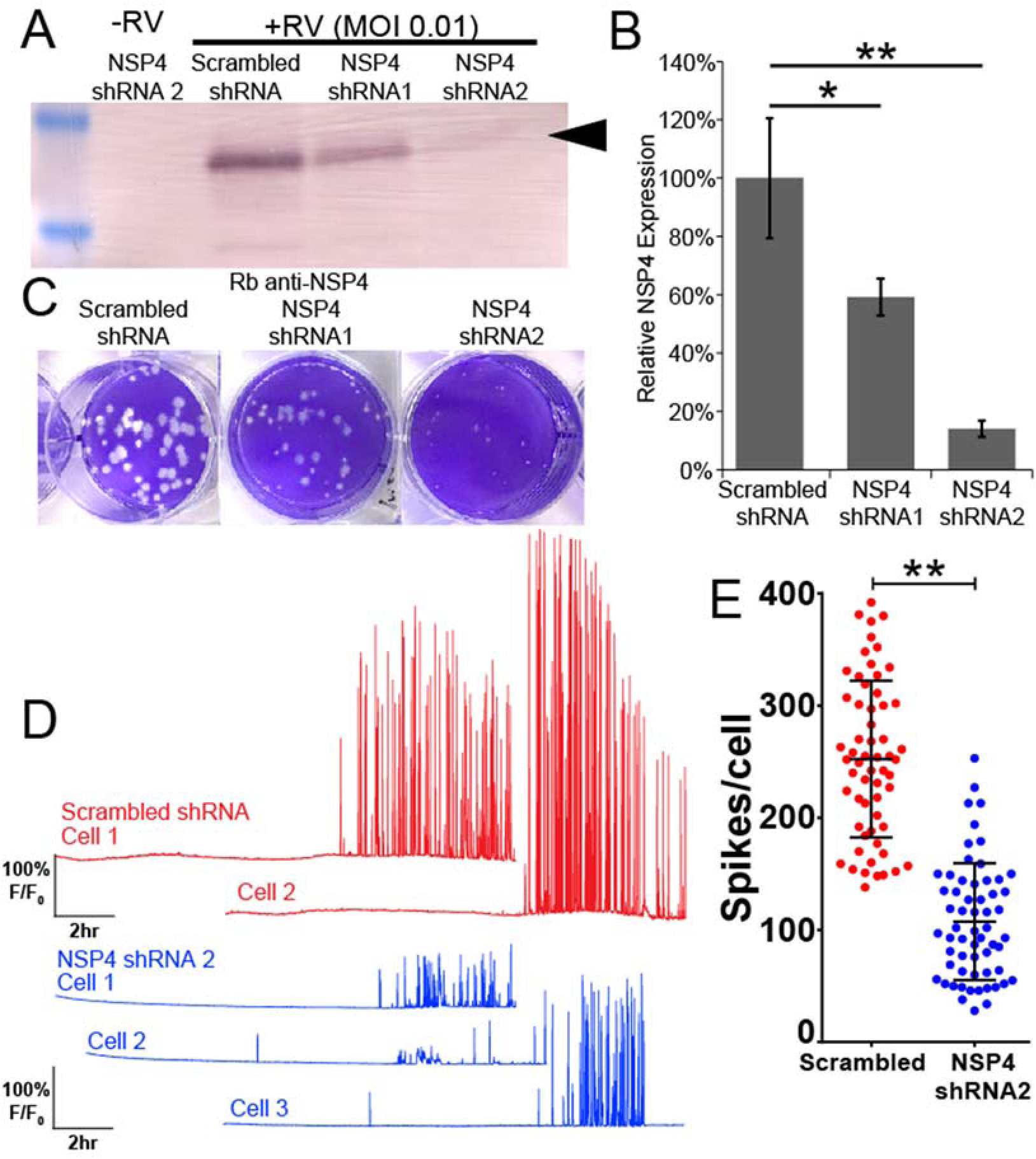
Knockdown of NSP4 reduces the dynamic Ca^2+^ signaling. **(A)** Western blot for NSP4 expression in mock infected or RV-infected MA104-GCaMP6s/shRNA cell lines. Cells expressing either scrambled, NSP4 shRNA1, or NSP4 shRNA2 are as indicated. Arrow denotes full-length, glycosylated NSP4. **(B)** Densitometry analysis of western blots for NSP4 levels normalized to that of GAPDH levels, expressed as relative to NSP4 expressed in MA104-GCaMP6s/scrambled shRNA cells. Data shown are mean ± SD of 3 infections/condition and representative of 3 independent experiments. **(C)** Plaque assay of MA104-GCaMP6s/shRNA cell lines inoculated with SA114F (10^−6^ dilution). **(D)** Representative single-cell traces of relative GCaMP6s fluorescence (F/F_0_) from SA114F infection of cells expressing scrambled shRNA (red) or NSP4 shRNA2 (blue). **(E)** Number of Ca^2+^ spikes (F/F_0_ > 5%) from mock or RV-infected cells inoculated with MOI0.1 SA114F. Data shown as mean ± SD of 60 cells/condition. **p<0.01.

### RV-induced Ca^2+^ spikes require extracellular and ER Ca^2+^ pools

NSP4 elevates cytosolic Ca^2+^ by activating both uptake of extracellular Ca^2+^ and release of ER Ca^2+^ pools. Thus, we next sought to characterize which pools of Ca^2+^ were critical for supporting the RV-induced Ca^2+^ spikes. First, we tested whether extracellular Ca^2+^ influenced the RV-induced Ca^2+^ spikes. We infected MA104-GCaMP5G cells with SA114F (MOI 1), and at 1 HPI replaced the media with either normal media containing 2 mM Ca^2+^, media without Ca^2+^ (supplemented with 1.8 mM EDTA), or media supplemented with Ca^2+^ for a 10 mM final concentration. The imaging data shows that decreasing extracellular Ca^2+^ strongly reduced the number and duration of Ca^2+^ signaling (Fig. 6A, light blue), whereas cells maintained in media with 2 mM or 10 mM extracellular Ca^2+^ showed increased dynamic Ca^2+^ signaling (Fig. 6A, red & purple). As before, mock-infected cells in each condition exhibited little to no induction of the Ca^2+^ signaling (Fig. 6A, black lines). Quantitation of the number of Ca^2+^ spikes showed that RV-infected cells in low extracellular Ca^2+^ exhibited significantly fewer Ca^2+^ spikes than that of cells in normal extracellular Ca^2+^, but this was still greater than that of mock-infected cells (Fig. 6B). Interestingly, there was no difference in the number of Ca^2+^ spikes per cell when maintained in 2 mM versus 10 mM Ca^2+^ media (Fig. 6B). However, the traces indicated that the magnitude of the Ca^2+^ spikes were greater in high Ca^2+^ media. Thus, we then determined the Ca^2+^ spike amplitude for the top 50 Ca^2+^ spikes, and while this was highly variable from cell-to-cell, this trended to be greater with higher extracellular Ca^2+^ concentrations (Fig. 6C). Together, these data indicate that normal extracellular Ca^2+^ levels are critical for the RV-induced Ca^2+^ spikes, which could occur both through discrete Ca^2+^ influx events through the plasma membrane, and by influx of extracellular Ca^2+^ serving to maintain ER Ca^2+^ stores to feed ER Ca^2+^ release events.

**Figure 6.**
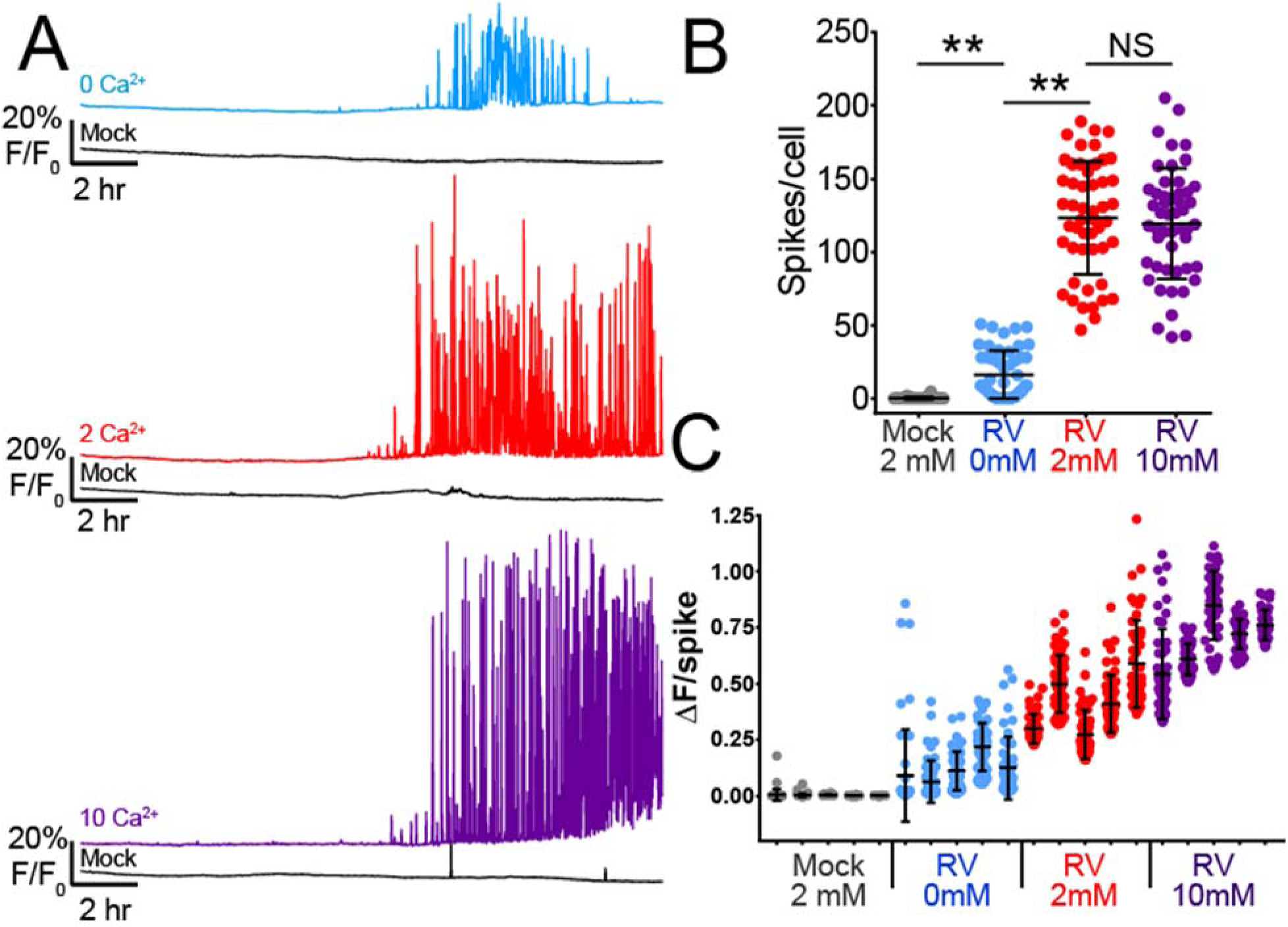
Ca^2+^ signaling requires extracellular Ca^2+^. **(A)** Representative single-cell traces of relative GCaMP5G fluorescence (F/F_0_) from mock (black lines) or SA114F infected cells in Ca^2+^-free media (0Ca^2+^, light blue), normal media (2Ca^2+^, red), and high Ca^2+^ media (10Ca^2+^, purple). **(B)** Number of Ca^2+^ spikes (F/F_0_ > 5%) from mock or RV-infected cells inoculated with MOI 1 SA114F and maintained in the indicated media. Data shown as mean ± SD of 50 cells/condition. **p<0.01. **(C)** Ca^2+^ spike amplitude of the top 50 Ca^2+^ spikes from five representative mock and RV-infected cells in each media condition.

The ER is the major intracellular Ca^2+^ store, and RV NSP4 has been shown to decrease ER Ca^2+^ levels both during infection and by recombinant expression ^6, 7^. However, controversy remains about whether RV causes a sustained depletion in ER Ca^2+^ ^22^. Thus to directly characterize the change in ER Ca^2+^ during RV infection, and determine how this relates to the dynamic cytosolic Ca^2+^ spikes, we generated an MA104 cell line co-expressing R-GECO1.2 and GCEPIAer (MA104-RGECO1/GCEPIAer), in which R-GECO1.2 is a red fluorescent cytoplasmic GECI and GCEPIAer is a green fluorescent ER-targeted GECI ^24, 41^. As above, infection with SA114F (MOI 1) induced highly dynamic cytoplasmic Ca^2+^ signaling by ∼8 hpi, as illustrated in two representative single-cell traces (Fig. 7A-B, red traces; Supplementary Video 6 online). Concomitant with the onset of the cytoplasmic Ca^2+^ signaling was an equally dynamic decrease of ER Ca^2+^ that persisted throughout the rest of the infection (Fig. 7A-B, green traces). We examined the relationship between the cytoplasmic Ca^2+^ spikes and ER Ca^2+^ troughs more closely from 8-12 hpi (Fig. 7B), which was during the onset of these signaling events. First, we found that the onset of Ca^2+^ signals in the cytoplasm coincided with the ER Ca^2+^ release events (Fig. 7B, black arrowheads). The persistent decrease in ER Ca^2+^ observed was driven primarily by this continuous signaling, such that the ER Ca^2+^ level never recovered to the baseline level (Fig. 7A). Interestingly, a small number of ER Ca^2+^ troughs were not associated with a concomitant cytoplasmic Ca^2+^ spike (Fig. 7B, magenta arrowheads). Over the course of the long-term imaging experiment, mock-infected cells exhibited a 10% decrease in GCEPIAer fluorescence, but RV-infected cell had a 30% decrease (Fig. 7C), which occurred rapidly from 8-12 HPI (Fig. 7A). The decrease in GCEPIAer in mock-infected cells likely represents modest photobleaching of GCEPIAer over the imaging experiment.

**Figure 7.**
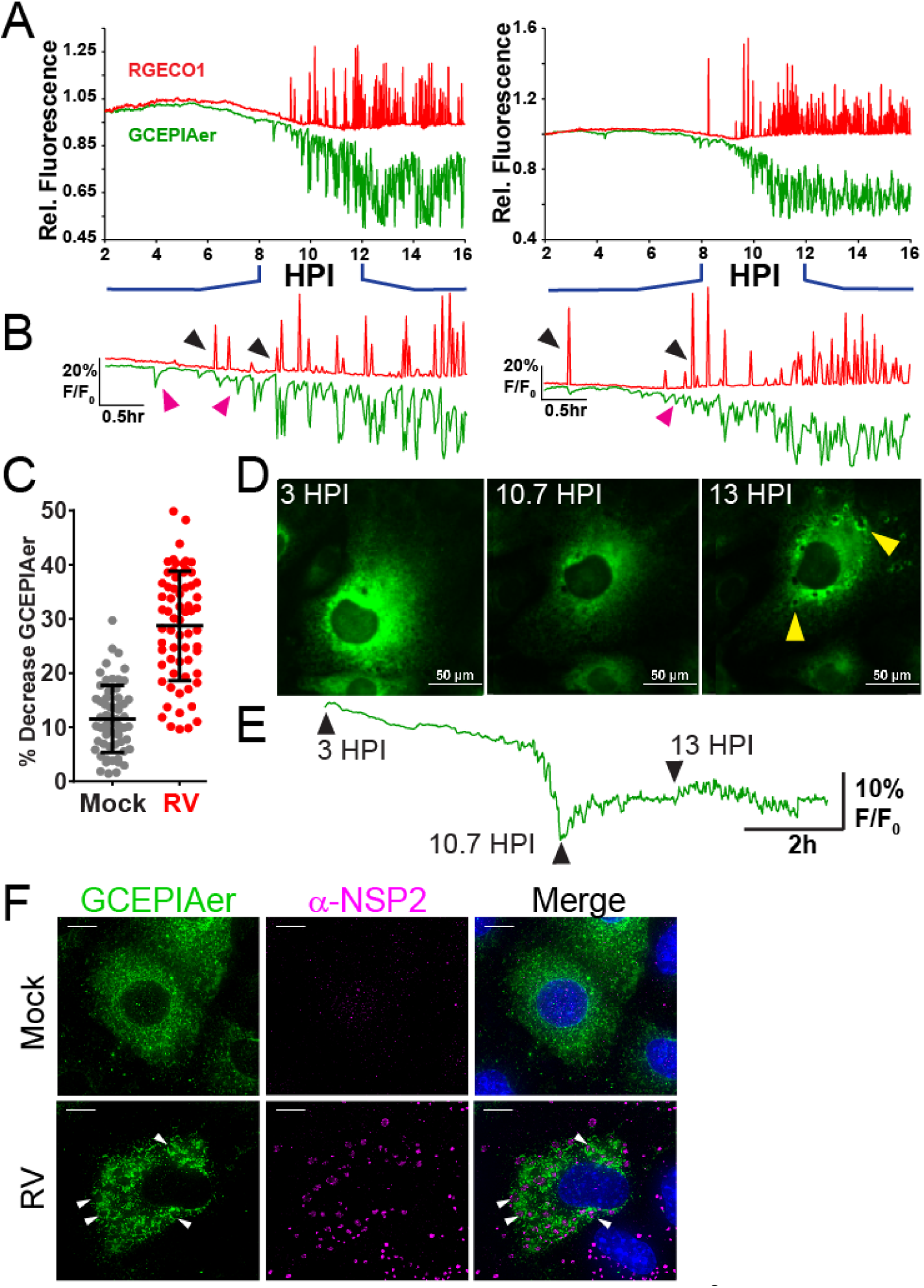
RV induces dynamic decreases in ER Ca^2+^ that coincide with cytoplasmic Ca^2+^ signals. **(A)** Two representative single-cell traces of relative RGECO1.2 (red) and GCEPIAer (green) fluorescence (F/F_0_, both RGECO1.2 and GCEPIAer are shown on the same scale) from MA104-RGECO/GCEPIAer cells infected with SA114F at MOI 1. **(B)** Traces from (A) expanded to show details from 8-12 HPI. Black arrowheads indicate ER Ca^2+^ troughs that correspond to cytoplasmic Ca^2+^ spikes. Magenta arrowheads indicate ER Ca^2+^ troughs that lack a corresponding cytoplasmic Ca^2+^ spike. **(C)** Percent decrease in ER Ca^2+^ levels measured by GCEPIAer fluorescence. **(D-E)** Images of a representative RV-infected MA104-GCEPIAer cell (D) taken at 3, 7.75, and 10 HPI and the corresponding trace from that cell with arrowheads corresponding the images above. Formation of viroplasm structures (yellow arrowheads) are observed subsequent to the decrease in ER Ca^2+^ levels. **(F)** Deconvolution microscopy of mock or RV-infected MA104-GCEPIAer cells (MOI 0.25, fixed 24 hpi) stained with α-NSP2 [Dylight 549 (pink)] to detect viroplasms (white arrowheads). (scale bar = 10 µm)

During these studies, we observed that the ER-localized GCEPIAer protein is also redistributed during the RV infection, which is illustrated for a single cell in Fig. 7D-E and Supplementary Video 7. At the beginning of the imaging run (3 hpi), the GCEPIAer signal was high and localized throughout the ER in a reticular pattern (Fig. 7D-E, left) but by 10.7 HPI, RV-induced Ca^2+^ signaling had decreased GCEPIAer fluorescence to its nadir (Fig. 7D-E, middle), representing a substantial decrease in ER Ca^2+^. Approximately 2 hrs after the initial decrease in ER Ca^2+^, the GCEPIAer began accumulating into circular domains that are likely the ER-derived compartment surrounding viroplasms (Fig. 7D-E, right). These structures become more pronounced through the late stages of the infection, ∼13 hpi (Fig. 7D, arrows). While the absolute onset of the ER Ca^2+^ release events was variable, the formation of viroplasm-associated membranes subsequent to the decrease in ER Ca^2+^ was a consistent pattern among RV-infected cells (Supplementary Video 7 online). Using immunostaining and deconvolution microscopy, we confirmed that the structures are viroplasms because they contain RV nonstructural protein 2 (NSP2), a major component of viroplasms (Fig. 7F). Interestingly, during late stages of infection when viroplasms are forming, we detected a modest recovery in ER Ca^2+^ from its nadir (Fig. 7E), which may reflect the increased ^45^Ca^2+^ uptake previously observed^22^.

### SOCE blockers reduce RV-induced Ca^2+^ spikes

Since removing extracellular Ca^2+^ diminishes the RV-induced Ca^2+^ spikes (Fig. 6), cellular Ca^2+^ influx pathways are critical for these Ca^2+^ signals. Several host Ca^2+^ channels have been implicated in mediating Ca^2+^ entry into RV-infected cells, including SOCE channels, voltage-activated Ca^2+^ channels (VACC), and the sodium-calcium exchanger (NCX) ^10–12^. To determine which pathway(s) were important for the dynamic Ca^2+^ spikes in RV infection, we used pharmacological blockers targeting each pathway (2-APB for SOCE; D600 for VACC; KB-R7943 for NCX). MA104-GCaMP5G cells were infected with SA114F (MOI 1), and then treated with different concentrations of the blockers at 1 hpi and imaged to measure GCaMP5G fluorescence. None of the blockers exhibited cytotoxic effects to uninfected cells (data not shown). Cells treated with DMSO as a vehicle control exhibited the dynamic Ca^2+^ spikes as above (Fig. 8A, red trace). In contrast, cells treated with the SOCE blocker 2-APB exhibited a dose-dependent decrease in both the number and amplitude of the Ca^2+^ signaling (Fig. 8A, green traces). Traces from cells treated with the NCX blocker KB-R7943 showed a modest decrease in Ca^2+^ signaling (Fig. 8A, brown traces), whereas there was no difference in Ca^2+^ signaling for cells treated with D600 (data not shown). We also noted that RV-infected cells treated with 10 μM KB-R7943 underwent cell death more frequently than any other treatment, which was marked by a rapid increase in cytosolic Ca^2+^ and then lysis (Fig. 8A, arrowhead); however, cell death was not observed in uninfected cells treated with KB-R7943 (data not shown). We quantitated the number of Ca^2+^ spikes per cell, which showed a significant, dose-dependent decrease in the number of Ca^2+^ spikes for both 2-APB-treated (Fig. 8B, green) and, to a lesser extent, KB-R7943-treated (Fig. 8B, brown) cells, but no difference for D600-treated cells (Fig. 8B, blue). We further investigated the effects of 2-APB and KB-R7934 by examining the amplitude of the largest 50 Ca^2+^ spikes of three representative cells shown in Fig. 8C-D. Treatment with 2-APB showed a dose-dependent decrease in the Ca^2+^ spike amplitude (Fig. 8C), consistent with the single-cell traces, but treatment with KB-R7943 showed no difference in spike amplitude.

**Figure 8.**
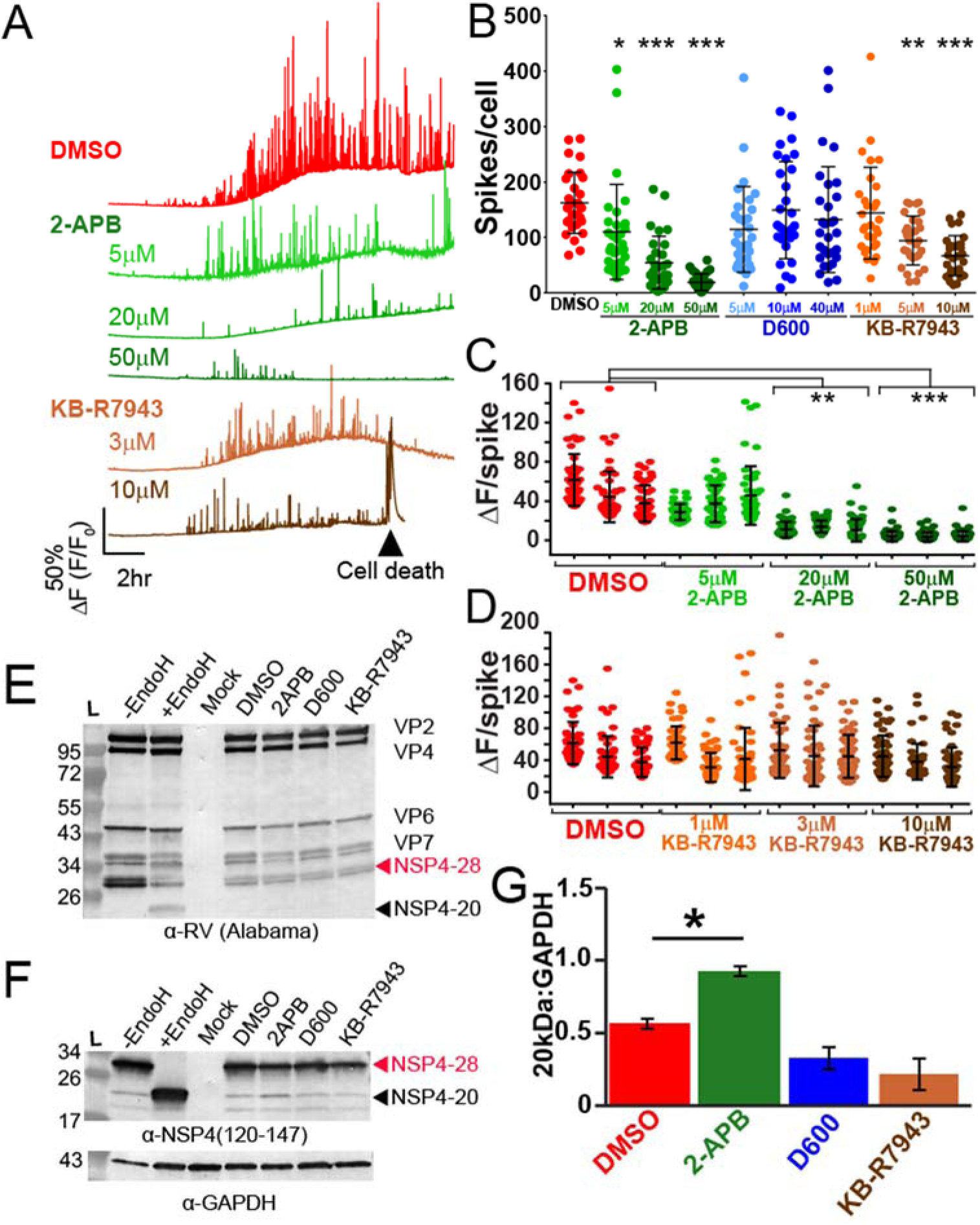
RV-induced Ca^2+^ signaling is blocked by the SOCE blocker 2APB. **(A)** Representative single-cell traces from MA104-GCaMP5G cells infected with SA114F MOI 1 and treated with DMSO vehicle alone or the indicated doses of 2APB or KB-R7943. Cell death of RV-infected cells treatment with 10 uM KB-R7943 was frequently observed (black arrowhead). **(B)** Number of Ca^2+^ spikes (F/F_0_ > 5%) from RV-infected cells inoculated with MOI 1 and treated with the indicated concentration of 2APB, D600 or KB-R7943. **(C-D)** Ca^2+^ spike amplitude of the top 50 Ca^2+^ spikes from three representative RV-infected cells treated DMSO (vehicle) or the indicated concentration of 2APB (C) or KB-R7943 (D). **(E-F)** Western blot analysis of MA104-GCaMP5G cells mock or RV-infected MOI 1 and treated with DMSO (vehicle), 50 μM 2APB, 10 μM D600, or 10 μM KB-R7943. Control RV-infected lysates treated with Endoglycosadase H (+EndoH) or untreated (-EndoH) are also shown. Blots were detected with α-RV (E) or α-NSP4(120-147) (F) and α-GAPDH (F, bottom) was the loading control for both. **(G)** ImageJ analysis of unglycosylated NSP4 (20kDa) normalized to GAPDH from RV-infected cells treated with the different blockers. Data are the mean ± SD of three independent infections per condition. *p<0.05

Since elevated cytosolic Ca^2+^ is critical for RV replication, we examined RV protein levels by immunoblot to determine whether 2-APB or KB-R7943 reduced the Ca^2+^ signaling by merely blocking RV or NSP4 protein synthesis or protein stability^42^. Immunoblot detection with an anti-RV antisera (Fig. 8E) or an anti-NSP4 specific antisera (Fig. 8F) show that none of the Ca^2+^ channel blockers caused substantial decrease in RV or NSP4 protein levels. However, we observed that 2-APB treatment significantly increased the 20 kDa unglycosylated NSP4 band (Fig. 8F) by gel densitometry analysis (Fig. 8G).

Overall the SOCE blocker 2-APB was the most potent inhibitor of the RV-induced dynamic Ca^2+^ signaling, so we examined the effect of other SOCE blockers that also target the Orai1 Ca^2+^ channel. MA104 cells express the Orai1 Ca^2+^ channel and the STIM1 and STIM2 ER Ca^2+^ sensors, which are the core machinery for the SOCE pathway (Fig. 9A). While MA104 cells also express the Orai3 Ca^2+^ channel, this isoform is not activated by ER Ca^2+^ store depletion but arachidonic acid and leukotrienes ^43^. We tested four SOCE blockers (2-APB, BTP2, Synta66, and GSK7975A) for the ability to block thapsigargin-induced SOCE and found that all of them showed a similar inhibition of Ca^2+^ entry after ER store depletion (Fig. 9B). Thus, we treated SA114F-infected MA104-GCaMP5G cells with each of these blockers at ∼1 hpi and performed Ca^2+^ imaging to measure the RV-induced Ca^2+^ spikes (Fig. 9C-D). Representative single-cell traces illustrate that all the SOCE blockers inhibited the RV-induced dynamic Ca^2+^ signaling (Fig. 9C) and significantly inhibited the number of Ca^2+^ spikes per cell (Fig. 9D). The blockers displayed varying degrees of potency but 2-APB and BTP2 treatment caused the greatest decrease in RV-induced Ca^2+^ signaling (Supplementary Video 8 online). Further, we found that treatment with the SOCE blockers significantly reduced RV yield from MA104 cells (Fig. 9E), which is consistent with the importance of elevated cytosolic Ca^2+^ for RV replication^29^. As with the other Ca^2+^ channel blockers, we examined whether the SOCE blockers affected viral protein levels by immunoblot. As above, we found that 2-APB treatment increased the abundance of the 20 kDa unglycosylated NSP4 band (NSP4-20), and Synta66 treatment also caused a modest increase in NSP4-20 (Fig. 9F). In contrast to the other SOCE blockers, BTP2 treatment caused an overall decrease in RV proteins, which correlates with the strong suppression of RV-induced Ca^2+^ signaling during the infection (Fig. 9D & 9F). Together, these data support the previous observation that shRNA knockdown of the Ca^2+^ sensor STIM1 reduces RV replication, and further show that Ca^2+^ influx *via* SOCE channels is critical for RV-induced Ca^2+^ signaling and replication^10^.

**Figure 9.**
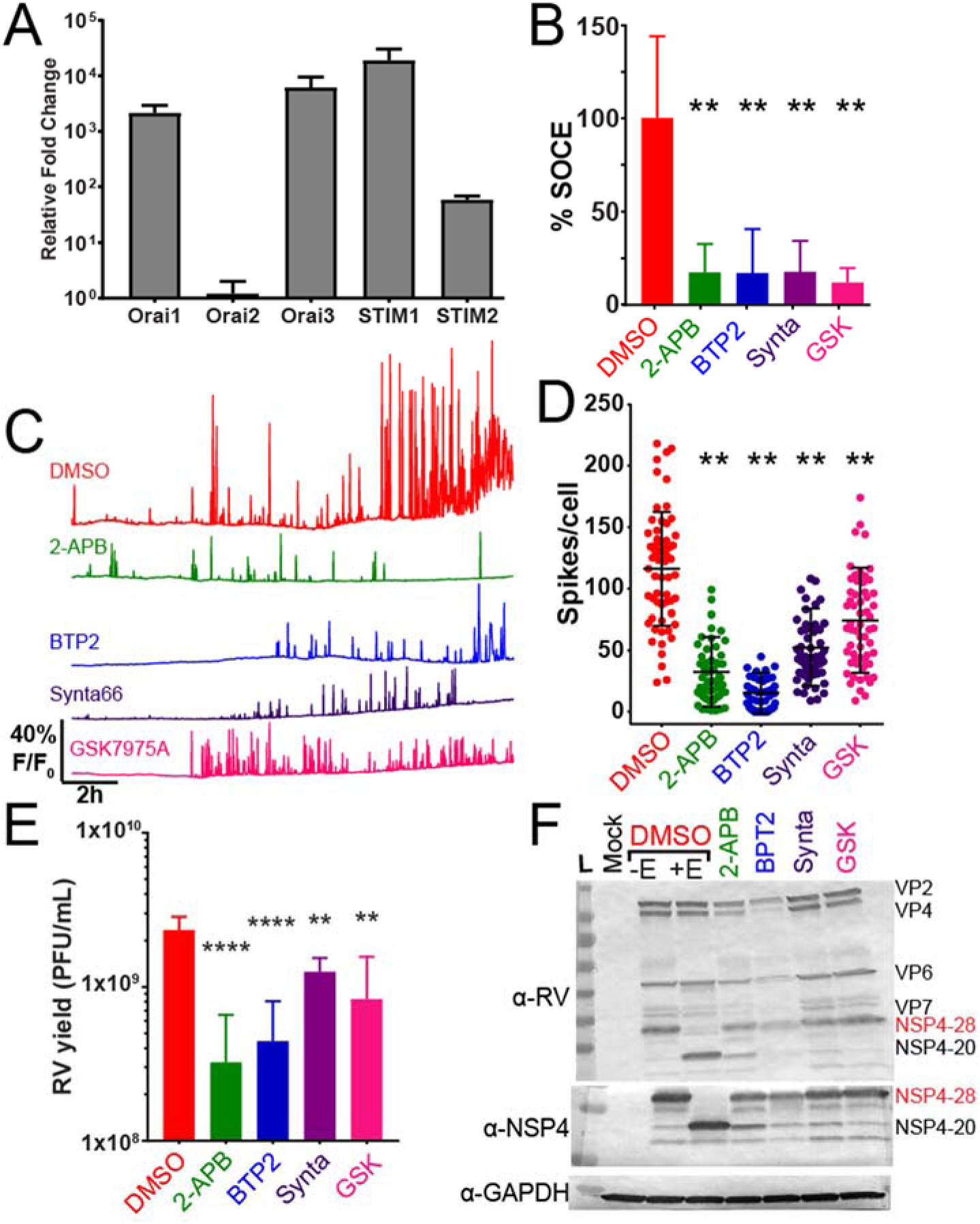
SOCE blockers reduce RV-induced Ca^2+^ signaling. **(A)** Relative mRNA expression of Orai1-3 and STIM1-2 genes in MA104 cells. Expression is normalized to 16S rRNA and graphed relative to Orai2. **(B)** SOCE was activated by treatment with 0.5 μM thapsigargin in Ca^2+^-free buffer and the amount of SOCE relative to DMSO-alone (vehicle) for different SOCE blockers determined. Data are the mean ± SD of three independent runs. **p<0.01. **(C)** Representative single-cell traces from MA104-GCaMP5G cells infected with SA114F MOI 1 and treated with DMSO vehicle alone or the indicated doses of 50 μM 2APB, 10 μM BTP2, 10 μM Synta66, or 10 μM GSK7975A. **(D)** Number of Ca^2+^ spikes (F/F_0_ > 5%) from RV-infected cells inoculated with MOI 1 and treated with DMSO alone or the SOCE blockers. Data are the mean ± SD 60 cells/condition. **p<0.01 by one-way ANOVA. **(E)** SA114F yield from MA104-GCaMP5G cells treated with DMSO or the SOCE blockers. Data are the mean ± SD of three independent infections. ****p<0.0001; **p<0.01 by one-way ANOVA. **(F)** Western blot analysis of MA104-GCaMP5G cells mock or RV-infected MOI 1 and treated with DMSO or the SOCE blockers. Control RV-infected lysates treated with Endoglycosadase H (+EndoH) or untreated (- EndoH) are also shown. Blots were detected with α-RV, α-NSP4(120-147), and α-GAPDH for the loading control.

### Human intestinal enteroid characterization of RV-induced Ca^2+^ signaling

Although MA104 cells provide a robust model for RV replication and form a single epithelial sheet ideal for microscopy studies, they are neither of human nor of intestinal cell origin. Human intestinal enteroids (HIEs) have been developed as a model *in vitro* system of the epithelial cells of the small intestine, and support RV infection and replication, particularly for human RV strains ^26, 44^. HIEs are grown in “mini-gut” three-dimensional (3D) cultures from human intestinal stem cells and are non-transformed cells, which make them a biologically relevant system to study the GI epithelium^45^. Thus, we sought to determine if the dynamic cytosolic Ca^2+^ signaling observed in MA104 cells were also observed in HIEs with RV infection.

We created jejunum HIEs stably expressing the green cytoplasmic GECI GCaMP6s (jHIE-GCaMP6s) using lentivirus transduction. To test the response of GCaMP6s to cytoplasmic Ca^2+^ in the enteroids, we treated 3D jHIE-GCaMP6s stabilized in a diluted Matrigel, with carbachol, a known Ca^2+^ agonist. Carbachol treatment of jHIE-GCaMP6s significantly increased GCaMP6s fluorescence 200-300% over the mock-treated jHIE-GCaMP6s (Fig. 10A-C). Thus, jHIE-GCaMP6s enteroids functionally report changes in cytoplasmic Ca^2+^ and can be used to examine RV-induced Ca^2+^ signaling.

**Figure 10.**
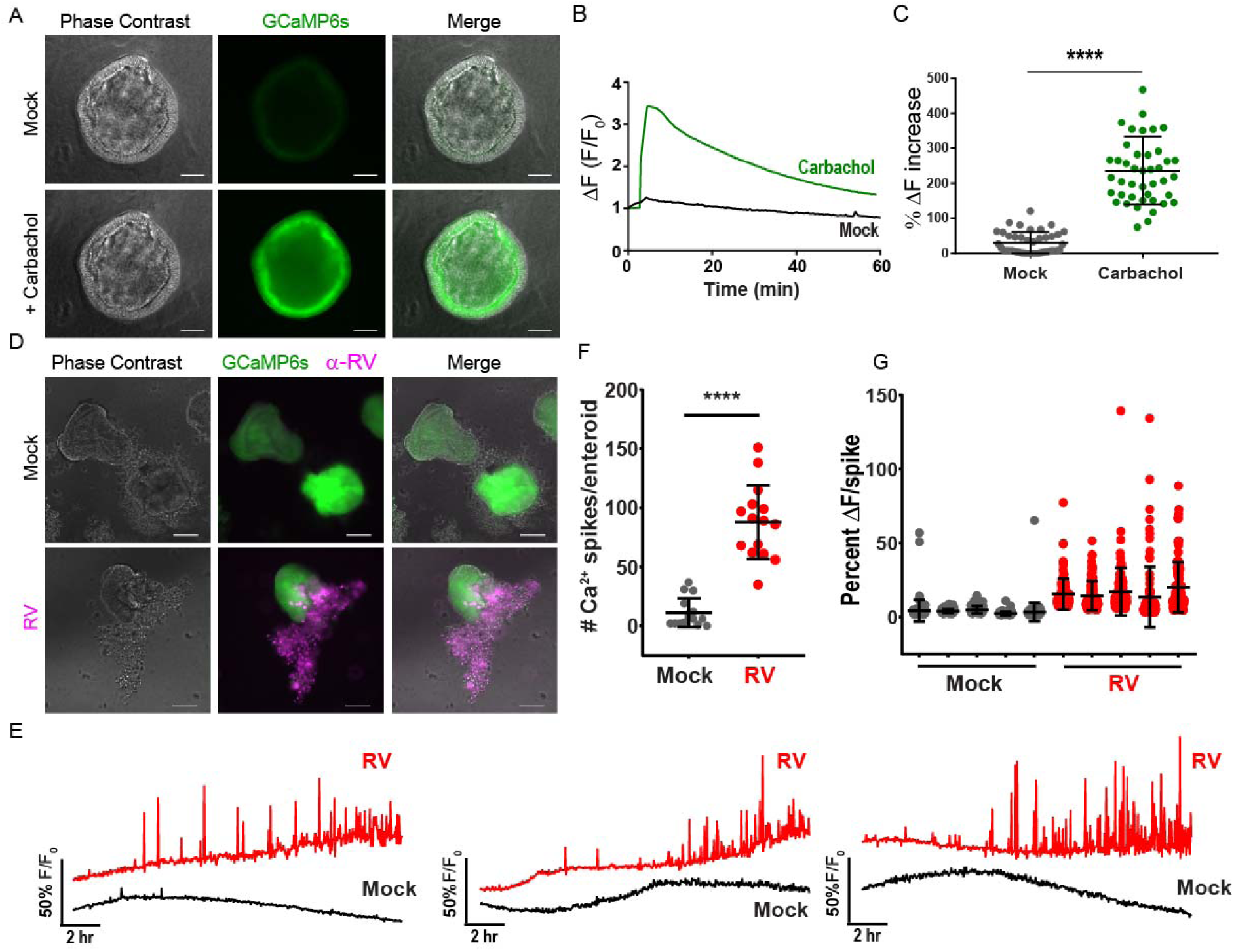
jHIE-GCaMP6s enteroids exhibit dynamic Ca^2+^ signaling during RV infection. **(A)** Representative images of jejunum human intestinal enteroids stably expressing GCaMP6s (jHIE-GCaMP6s) one minute after addition of 200 µM carbachol or media alone (mock). **(B)** Representative trace of jHIE-GCaMP6s treated with 200 µM carbachol or media alone. GCaMP6s fluorescence (F) was normalized to the baseline fluorescence (F_0_). **(C)** The maximum % increase in normalized fluorescence after addition of media (mock, N=41) or 200 µM carbachol (N=43), data combined from 3 independent experiments. **(D)** Representative immunofluorescence images of mock and Ito RV-infected jHIE-GCaMP6s (green) fixed at ∼24 hpi and stained for RV (pink). **(E)** Representative Ca^2+^ traces of whole jHIE-GCaMP6s enteroids either mock or Ito RV-infected between 6-23 hpi. GCaMP6s fluorescence (F) was normalized to the baseline fluorescence (F_0_). **(F)** Number of Ca^2+^ spikes in mock (N=15) or Ito RV-infected (N=15) HIEs and **(G)** the fold change in fluorescence for randomly selected 5 HIEs mock- and RV-infected. Data are representative of one experiment that was performed in triplicate. ****p<0.0001, scale bars = 100 µm.

We next tested if 3D jHIE-GCaMP6s enteroids would exhibit similar Ca^2+^ dynamics during RV infection as observed in MA104-GCaMP5G cells. jHIE-GCaMP6s enteroids were mock- or RV-infected with the human RV strain Ito, seeded into chamber slides in diluted Matrigel, and imaged every 2-3 minutes for phase contrast and GCaMP6s fluorescence throughout the RV infection (∼16 hrs). At 24 hpi, the HIEs were fixed and immunostained for RV antigens to confirm successful infection, which is evident by both infected cells within the HIEs as well as strong positive staining of the dead cells sloughed from the HIEs (Fig. 10D). Examination of the Ca^2+^ signaling showed little Ca^2+^ signaling activity in the mock-infected jHIE-GCaMP6s enteroids, but RV-infected enteroids exhibited significantly increased Ca^2+^ dynamics, as illustrated in representative traces from three mock- or RV-infected HIEs (Fig. 10E and Supplementary Video 9 online). Similar to the Ca^2+^ signaling observed in MA104 cells, initially there were no or only modest changes in cytosolic Ca^2+^, and the onset of strong and dynamic Ca^2+^ signals occurred ∼8-10 HPI. For HIEs it was not possible to accurately measure Ca^2+^ signaling at the single-cell level. We were able to track and measure Ca^2+^ signaling over the entire jHIE-GCaMP6s enteroid and quantify these changes as Ca^2+^ spikes/enteroid. We found that RV significantly increased the number of Ca^2+^ spikes/enteroid (Fig. 10F) and that the Ca^2+^ spike amplitudes are also substantially greater in RV-infected than in mock-infected jHIE-GCaMP6s enteroids (Fig. 10G). Thus, the RV-induced Ca^2+^ signaling in enteroids closely parallels that observed in MA104 cells and demonstrate that these dynamic Ca^2+^ signals are a biologically relevant aspect of how RV disrupts host Ca^2+^ homeostasis.

Since SOCE played a prominent role in the RV-induced dynamic Ca^2+^ signaling in MA104 cells, we investigated whether it was also critical for the Ca^2+^ signaling observed in HIEs. Similar to MA104 cells, jejunum-derived HIEs expressed the core SOCE proteins Orai1, STIM1, and STIM2, as well as the non-store operated Orai3 channel (Fig. 11A). The expression levels were not substantially altered by differentiating the jHIEs through removal of growth factors. To test whether SOCE is important for RV-induced Ca^2+^ signaling, we first tested 2-APB treatment of 3D jHIE-GCaMP6s enteroids either mock- or RV-infected with strain Ito. While RV infection increased the number of Ca^2+^ spikes per enteroid consistent with above (Fig. 11B), 2-APB treatment did not attenuate the Ca^2+^ signaling (Fig.11B-C). We speculated that the 3D format or the Matrigel used to support 3D HIEs might interfere with 2-APB blocking SOCE, so we repeated these studies using jHIE-GCaMP6s monolayers. First, we confirmed that 2-APB can block thapsigargin-induced SOCE in jHIE-GCaMP6s monolayers, which exhibited a 32% reduction in Ca^2+^ re-entry after store depletion (Fig. 11D). Interestingly, 2-APB shows a much less potent block of SOCE in enteroids than in MA104 cells, which exhibited a >80% inhibition of Ca^2+^ re-entry after store depletion (Fig. 9C). We also tested if VACC or NCX may contribute to RV-induced Ca^2+^ signaling in enteroids, but treatment with D600 or KB-R7943 did not reduce Ca^2+^ spikes in RV-infected jHIE-GCaMP6s monolayers (Fig. 11E). Nevertheless, 2-APB treatment of both mock-inoculated (Fig. 11F, black vs. grey traces) and RV-infected jHIE-GCaMP6s monolayers (Fig. 11F, red vs. blue traces) reduced the observed Ca^2+^ signaling, as illustrated in the representative traces (see Supplementary Video 10 online). We quantitated the Ca^2+^ signaling per FOV and confirmed that 2-APB treatment significantly reduced the number of Ca^2+^ spikes for both mock and RV-infected enteroids (Fig. 11G), as well as substantially reducing the amplitude of the Ca^2+^ signals (Fig. 11H). Thus, like the MA104 model, SOCE is critical for supporting the dynamic Ca^2+^ signaling induced in RV-infected jHIEs.

**Figure 11.**
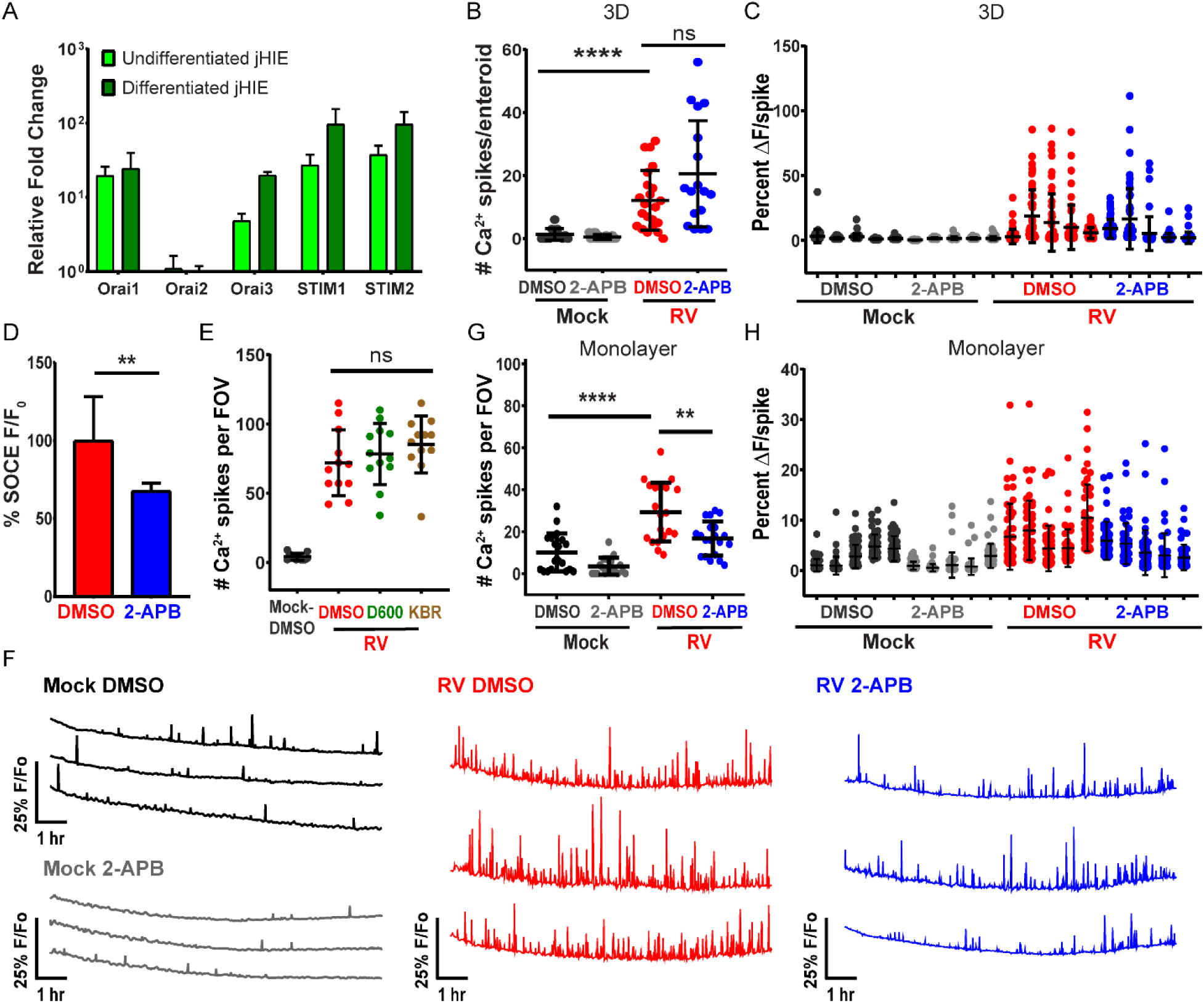
Blocking SOCE reduces Ca^2+^ signaling during RV infection in jHIE-GCaMP6s monolayers. **(A)** qPCR of Orai and stromal interaction molecule (STIM) mRNA transcripts normalized to 18S mRNA transcripts and fold change relative to Orai2 mRNA transcript levels in jejunum human intestinal enteroids (jHIEs), log scale (n = 3 biological replicates). **(B)** Number of Ca^2+^ spikes in 3D jHIE-GCaMP6s mock- or RV-infected and treated with DMSO (N=12, 18) or 50 µM 2APB (N=22, 17) and **(C)** the percent change in fluorescence for the highest 50 Ca^2+^ spikes for 5 HIEs of each condition for 6-18 hpi. GCaMP6s fluorescence (F) was normalized to the baseline fluorescence (F_0_). Data are representative of one experiment that was performed in triplicate. **(D)** Number of Ca^2+^ spikes per field-of-view (FOV) in mock- or RV-infected jHIE-GCaMP6s monolayers treated with DMSO, 10 µM D600, or 10 µM KB-R7943 between 8-22 hpi, data combined from 3 experiments. **(E)** Relative fluorescence increase (F/F_0_) due to store-operated calcium entry in jHIE-GCaMP6s monolayers after store depletion with 500 nM thapsigargin. Data combined from ≥6 experiments. **(F)** Representative Ca^2+^ traces per FOV of monolayers either mock- or RV-infected and treated with DMSO or 50 µM 2APB between 8-19 hpi. **(G)** Number of Ca^2+^ spikes per FOV and **(H)** the percent change in fluorescence for the highest 50 Ca^2+^ spikes for 5 FOVs of each condition for 8-19 hpi. Data combined from 5 experiments. (**p<0.01, ****p<0.0001)

## Discussion

A hallmark of RV infection, and several other viruses, is an elevation in cytosolic Ca^2+^ and decrease in ER Ca^2+^ stores, which facilitates virus replication and contributes to pathogenesis through a variety of downstream pathways ^1, 46^. The importance of RV-induced dysregulation of Ca^2+^ levels for many of these downstream pathways has been determined, but thus far characteristics of the Ca^2+^ signaling itself have not been extensively investigated^1, 5^. Thus, the primary goal of this study was to determine the nature of the RV-induced elevation in cytosolic Ca^2+^ and characterize how the dysregulation of Ca^2+^ signaling manifests during the infection. By leveraging GECI-expressing cell lines to perform long-term Ca^2+^ imaging, we found that RV induces a vast increase in Ca^2+^ signaling events that increased in frequency and magnitude over the course of the infection. These results are consistent with previous measurements of cytosolic Ca^2+^ in RV-infected cells that show a monophasic increase over time, which is similar to our imaging data when it is averaged out across the whole FOV (*i.e.*, a cell population). Yet, what is paradigm changing is that at the individual cell level, RV does not merely cause a steady increase in cytosolic Ca^2+^, but rather activates a cacophony of discrete Ca^2+^ signaling events. Further, by generating GECI-expressing HIEs, this study is the first characterization of RV-mediated Ca^2+^ signaling in normal, human small intestinal enterocytes. We found that the prominence of the Ca^2+^ spikes in RV-infected HIEs is similar to that in MA104 cells, underlining that this is a biologically relevant phenomenon.

The characterization of the RV-induced increase in cytosolic Ca^2+^ as a series of discrete, transient Ca^2+^ signals is an important new insight into the cellular pathophysiology of RV infection. Transient increases in cytosolic Ca^2+^ serve as pro-survival signals by activating phosphoinositide 3-kinase (PI3K) and by calcineurin-dependent NFAT activation. Further, Ca^2+^ oscillations stimulate mitochondrial Ca^2+^ uptake that enhances ATP synthesis, and this increase in mitochondrial metabolism contributes to cell survival pathways. In contrast, strong sustained elevation of cytosolic Ca^2+^ drives pro-apoptotic signaling through mitochondrial Ca^2+^ overload^20^. Thus, even though the mean cytosolic Ca^2+^ level is progressively increasing in the RV-infected cell, early activation of the intrinsic apoptotic cascade may be prevented because it occurs as hundreds of transient Ca^2+^ signals over hours. This premise is consistent with studies showing that early activation of PI3K during RV infection delays apoptosis^47^.

Concomitantly, the elevated Ca^2+^ signaling activates cellular pathways, such as autophagy, that promote RV replication and assembly of progeny virus^29^. Whether the initial Ca^2+^ dynamics enhance mitochondria bioenergetics or ATP synthesis early during RV infection has not been studied, but a loss of mitochondria membrane potential and decrease in ATP output occur at late stages of infection^47, 48^. Ultimately the massive increase in Ca^2+^ signaling damages the cell and triggers cell death, and cell lysis was observed in our time-lapse imaging, but this data cannot differentiate whether this was through apoptosis, necrosis, and/or pyroptosis^18, 47, 49^. Thus, RV exploitation of discrete Ca^2+^ signals, rather than a sustained increase in cytosolic Ca^2+^, may function in concert with other RV anti-apoptotic proteins, such as NSP1, to forestall the onset of cell death and enable sustained viral replication.

Due to GECI photostability, we were, for the first time, able to perform Ca^2+^ imaging of individual cells throughout the RV infection. The increased Ca^2+^ spikes are the predominant feature of this imaging, and the SA11-mRuby reporter virus demonstrates that the onset of the increased Ca^2+^ signaling correlates with RV protein synthesis as well as input virus dose. Yet, despite individual cells exhibiting very distinct Ca^2+^ signaling traces, a general pattern emerges: [i] early in infection Ca^2+^ signaling remains at basal levels; [ii] onset of increased Ca^2+^ signaling is characterized by low-amplitude Ca^2+^ spikes relatively; and then [iii] very high-amplitude Ca^2+^ spikes become predominant, which results in elevated cytosolic Ca^2+^ levels. The progression of the RV-induced Ca^2+^ signals have important implications for characterizing how Ca^2+^-regulated cellular processes are influenced by RV infection. For example, RV induces autophagy through increased cytosolic Ca^2+^, thereby activating calcium/calmodulin-dependent kinase kinase-β (CaMKKβ)^29^. This raises several questions: *When* after onset of the aberrant Ca^2+^ signaling is CaMKKβ activated? *How many* Ca^2+^ signals are needed to induce autophagy? Similar questions can be asked about the role of these Ca^2+^ signals in RV-induced apoptosis, cytoskeletal rearrangement, and serotonin and chloride secretion^26, 47, 50, 51^. Tracking these dynamic relationships poses a challenge that may be addressed by further engineering GECI-expressing cell lines/HIEs to express other biosensors such that both processes can be measured throughout the infection.

Many studies show that RV infection (or NSP4 expression) reduces the ER Ca^2+^ stores based on a blunted cytosolic Ca^2+^ release in response to agonists (*e.g.*, ATP) or thapsigargin treatment to prevent SERCA-mediated refilling^22, 52^. However, other results show increased in radioactive ^45^Ca^2+^ loading into the ER in RV-infected cells, which is hypothesized to be due to an increase in Ca^2+^ binding proteins (*e.g.*, VP7 or ER chaperone proteins)^7, 11^. Thus, controversy remains about whether RV causes a decrease in the ER Ca^2+^ store. To address this question, we developed MA104-RGECO1.2/GCEPIAer cells to directly measure cytosolic and ER Ca^2+^ together during the RV infection. RV induces a dynamic decrease in ER Ca^2+^ levels that occurs in conjunction with the increase in cytosolic Ca^2+^ signaling. In most instances, the cytosolic Ca^2+^ spike correlated with a decrease in ER Ca^2+^, indicating release of ER Ca^2+^ substantially contributes to the increased cytosolic Ca^2+^ signaling. Further, despite the 30% reduction in steady-state ER Ca^2+^, the dynamic nature of the ER Ca^2+^ signaling suggests that SERCA pumps continually work to refill the ER. The observed depletion of ER Ca^2+^ levels is consistent with the blunted cytosolic response to Ca^2+^ agonists like ATP, the NSP4 function as a Ca^2+^-conducting viroporin in the ER, and the activation of the ER Ca^2+^ sensor STIM1^7, 9, 10^. In contrast, it is more difficult to reconcile the previously observed increase in ^45^Ca^2+^ loading into the ER with the 30% reduction in steady-state ER Ca^2+^ levels detected by GCEPIAer imaging in this study. It has been hypothesized that increased ^45^Ca^2+^ loading may be due to increased ER Ca^2+^ buffering capacity, caused by the high levels of RV VP7 and/or chaperones BiP and endoplasmin^11^. However, our data suggest this is unlikely to be the case because this would sequester Ca^2+^ and render GCEPIAer unresponsive to changes in ER Ca^2+^^24^, yet this is not the case because GCEPIAer remains dynamic throughout the infection. Alternatively, the increase ^45^Ca^2+^ may reflect loading into the ER-derived autophagy-like microdomains that surround viroplasms, which we observed form after the initial depletion in ER Ca^2+^ and during a partial recovery of ER stores^29, 53^. These ER microdomains are the site VP7 assembly onto nascent RV particles, which requires high Ca^2+^, so Ca^2+^ sequestration in these microdomains may occur independently of the rest of the ER. Future studies using GCEPIAer and viroplasm-targeted GECIs are needed to determine whether the ER and viroplasm-associated membranes are functionally distinct compartments.

The pleiotropic functions of NSP4 are responsible for the RV-mediated dysregulation of host Ca^2+^ homeostasis through the ion channel function of iNSP4 and Ca^2+^ agonist function of the secreted eNSP4 enterotoxin ^5, 9, 10^. Our data show that NSP4 governs the dynamic Ca^2+^ signaling induced by RV infection since NSP4 knockdown significantly abrogated the number and amplitude of the Ca^2+^ spikes. Unfortunately, it is not possible to determine the relative roles of iNSP4 versus eNSP4 in the induction of the Ca^2+^ spikes from these data because the shRNA decreased total NSP4 synthesis, and therefore both pathways would be attenuated. The importance of NSP4 for the Ca^2+^ signaling is also demonstrated by the extremely different Ca^2+^ signaling profiles of OSUa- and OSUv-infected cells. These differences correlate with the attenuated elevation of cytosolic Ca^2+^ caused by recombinant OSUa NSP4 both when expressed in Sf9 cells (*i.e.*, iNSP4) and exogenous treatment of cells (*i.e.*, eNSP4)^40^. The attenuated NSP4 phenotype is the result of mutations in the NSP4 enterotoxin domain, indicating that this domain is critical for induction of the Ca^2+^ spikes by OSU^25, 40^. However, it is important to note that these two viruses are not isogenic so the genetic backgrounds of the OSUa and OSUv NSP4 are different, requiring further Ca^2+^ imaging studies using recombinant RV bearing these attenuating NSP4 mutations to fully dissect the relative importance iNSP4- and eNSP4-mediated Ca^2+^ signaling.

NSP4 is the trigger of the dynamic Ca^2+^ signaling, yet these signals are maintained through host Ca^2+^ channels and signaling pathways both in the ER and PM. Removal of extracellular Ca^2+^ significantly attenuated the Ca^2+^ spikes, demonstrating that Ca^2+^ influx is crucial for these signals. Three classes of Ca^2+^ channels (SOCE, NCX, and VACC) have been implicated RV-induced Ca^2+^ influx^10–12, 23^. Our results using different pharmacological blockers indicate SOCE is the primary Ca^2+^ influx pathway that supports the RV-induced dynamic Ca^2+^ spikes, both in MA104 cells and in HIEs. Blocking SOCE significantly reduced the number and amplitude of the RV-induced Ca^2+^ spikes. However, the Orai1 SOCE channel is a very low conductance channel so Ca^2+^ entry through PM Orai1 is unlikely to generate the high amplitude Ca^2+^ spikes observed during RV infection. The nature of the Ca^2+^ spikes, and the fact that most of them coincide with ER Ca^2+^ troughs, indicates that the signals detected are ER Ca^2+^-release events. ER Ca^2+^ release could occur either through iNSP4 or activation of the IP^3^-Receptor Ca^2+^ channel, and SOCE serves to maintain these signals by sustaining ER Ca^2+^ refilling. Since elevated Ca^2+^ levels are critical for RV replication, the attenuated Ca^2+^ signaling caused by the SOCE blockers significantly reduced RV yield^29^. Interestingly, blocking SOCE in HIEs significantly reduced the Ca^2+^ spikes, but the effect was less pronounced than in MA104 cells, suggesting other pathway(s) may exist that support RV-induced Ca^2+^ spikes in HIEs.

In summary, RV dysregulates host Ca^2+^ homeostasis by a massive and progressive increase in discrete Ca^2+^ signaling events, mainly from ER Ca^2+^ release. Many viruses elevate cytosolic Ca^2+^ and alter ER Ca^2+^, leading us to question whether dynamic Ca^2+^ spikes, as seen in RV infection, is a common manifestation for virus-induced Ca^2+^ signaling. If so, the host channels that support these Ca^2+^ signals, such as Orai1, may represent novel targets for broadly acting host-directed antiviral therapeutics.

## Acknowledgments

We would like to thank Dr. Lennart Svensson and Dr. Marie Hagbom for sharing their stocks of the porcine OSUa and OSUv viruses. This work was supported in part by NIH grants K01DK093657, R03DK110270, R01DK115507 (PI: J. M. Hyser), and R01AI080656 and U19AI116497 (PI: M. K. Estes). Trainee support for A.C.G. was provided by NIH grants F30DK112563 (PI: A.Chang-Graham) and the BCM Medical Scientist Training Program and support for both A.C.G. and A.C.S was provided by the Integrative Molecular and Biomedical Sciences Graduate Program (T32GM008231, PI: D. Nelson). This project was supported in part by PHS grant P30DK056338, which supports the Texas Medical Center Digestive Diseases Center (TMC-DDC) Gastrointestinal Experimental Model Systems (GEMS) Core and the Cellular and Molecular Morphology Core. Funding support for the BCM Integrated Microscopy Core includes the NIH (DK56338, CA125123), CPRIT (RP150578, RP170719), the Dan L.

Duncan Comprehensive Cancer Center, and the John S. Dunn Gulf Coast Consortium for Chemical Genomics. We would like to thank Xi-Lei (Shelly) Zeng and Xiaomin Yu for their help with enteroid cultures and media, and Drs. Michael Mancini and Fabio Stossi for deconvolution microscopy assistance. FACS sorting of cell lines utilized the BCM Cytometry and Cell Sorting Core with funding from the CPRIT Core Facility Support Award (CPRIT-RP180672), the NIH (CA125123 and RR024574), and the expert assistance of Joel M. Sederstrom.

## Author Contributions

JH, ACG, JP, AS, JC and MKE designed the experiments and discussed the data. JH, ACG, JP, NR, and AS conducted the calcium imaging experiments and analyzed the data with JH and ACG. JC and MKE provided key reagents including the shRNA knockdown cells. JP conducted the western blot and plaque assays and analyzed data with JH. AS and ACG conducted and analyzed qPCR experiments. JH and ACG wrote the manuscript, and all authors contributed to revisions of the paper.

