## Supplemental video 1 for "Rotavirus Calcium Dysregulation Manifests as Dynamic Calcium Signaling in the Cytoplasm and Endoplasmic Reticulum"

### Supplementary Figure

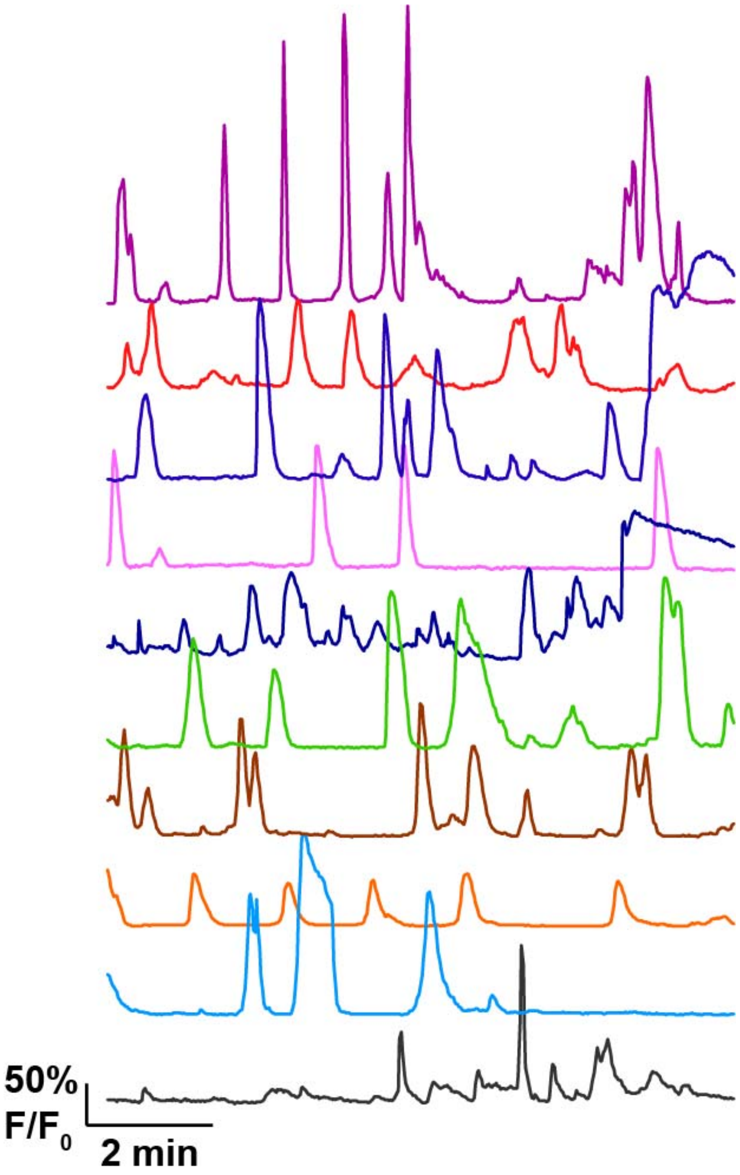


**Supplementary Figure 1.** Representative $Ca^{2+}$ traces ($F/F_0$) from 10 MA104-GCaMP5G cells infected with SA114F at MOI 10. Cells were imaged for 10 minutes at ~7 hpi with an image acquisition frequency of 1 image/1.5 seconds. These data correspond to Supplementary Video 3 online.
